# IKBKB reduces huntingtin aggregation by phosphorylating Serine 13 via a non-canonical IKK pathway

**DOI:** 10.1101/2022.12.05.519070

**Authors:** Cristina Cariulo, Paola Martufi, Margherita Verani, Leticia Toledo-Sherman, Ramee Lee, Celia Dominguez, Lara Petricca, Andrea Caricasole

## Abstract

N-terminal phosphorylation at residues T3 and S13 is believed to have important beneficial implications for the biological and pathological properties of mutant huntingtin, where IKBKB was identified as a candidate regulator of huntingtin N-terminal phosphorylation. The paucity of mechanistic information on IKK pathways, together with the lack of sensitive methods to quantify endogenous huntingtin phosphorylation, prevented detailed study of the role of IKBKB in Huntington’s disease. Using novel ultrasensitive assays, we demonstrate that IKBKB can regulate endogenous S13 huntingtin phosphorylation in a manner dependent on its kinase activity and known regulators. We found that the ability of IKBKB to phosphorylate endogenous huntingtin S13 is mediated through a non-canonical IRF3-mediated IKK-pathway, distinct from the established involvement of IKBKB in mutant huntingtin’s pathological mechanisms mediated via the canonical pathway. Furthermore, increased huntingtin S13 phosphorylation by IKBKB resulted in decreased aggregation of mutant huntingtin in cells, again dependent on its kinase activity. These findings point to a non-canonical IKK-pathway linking S13 huntingtin phosphorylation to the pathological properties of mutant huntingtin aggregation, thought to be significant to Huntington’s disease.

## Introduction

In Huntington’s disease (HD) an expansion of a CAG repeat within exon 1 of the *huntingtin* (*HTT*) gene, which produces a huntingtin (HTT) protein with an expanded polyglutamine (polyQ) repeat, leads to a progressive and fatal neurodegenerative pathology (Bates, Dorsey et al., 2015, McColgan & Tabrizi, 2018, Ross & Tabrizi, 2011). Mutant HTT fragments comprising the polyQ repeat, produced by alternative splicing and/or proteolytic cleavage, are widely believed to contribute significantly to HD through the propensity of such fragments to misfold and form aggregates (Bates et al., 2015, Landles, Sathasivam et al., 2010, Mangiarini, Sathasivam et al., 1996, Sathasivam, Neueder et al., 2013).

HD onset and progression are known to be influenced by a number of modifiers (Flower, Lomeikaite et al., 2019, Genetic Modifiers of Huntington’s Disease Consortium. Electronic address & Genetic Modifiers of Huntington’s Disease, 2019, Goold, Flower et al., 2019, Holmans, Massey et al., 2017, Lee, Chao et al., 2017), some of which may influence pathology by facilitating CAG-repeat expansion in somatic cells. Other evidence indicates that disease modification may be attainable through mechanisms directly affecting mutant HTT. Hence, protein sequences flanking the polyQ repeat—such as the first 17 residues of the HTT protein (N17 domain) and the proline-rich domain at the N- and C-termini of the polyQ repeat, respectively—has been shown to profoundly alter the biological and biophysical properties of these mutant HTT fragments in isolated proteins, cells and of the full length mutant HTT protein in vivo (Chatterjee, Steffan et al., 2021, Saudou & Humbert, 2016). Since the N17 domain is the target of several post-translational modifications (PTMs; (Ehrnhoefer, Sutton et al., 2011, Saudou & Humbert, 2016)), significant attention has focussed on residues associated with PTMs that may be modulated pharmacologically by targeting relevant enzymes, with the aim of altering the pathological properties of mutant HTT. Indeed, studies with phosphor-mimetic or phosphor-abrogative substitutions of residues T3 and S13/S16 have revealed a key role for these residues in regulating mutant HTT properties in isolated proteins (Crick, Ruff et al., 2013, Kelley, Huang et al., 2009, Mishra, Hoop et al., 2012), cells (Aiken, Steffan et al., 2009, Branco-Santos, Herrera et al., 2017, Maiuri, Woloshansky et al., 2013, Thompson, Aiken et al., 2009, Zheng, Li et al., 2013), and mice (Gu, Cantle et al., 2015), where phosphor-mimetic mutants invariably mitigate the effects of polyQ expansion. More recently, the availability of HTT proteins bearing bona-fide phosphorylation at these residues has confirmed the key role for T3 and S13/S16 in modulating mutant HTT phenotypes in vitro (Cariulo, Azzollini et al., 2017, Chiki A., 2017, DeGuire, Ruggeri et al., 2018). Parallel advances in developing assays that can specifically detect and quantify HTT phosphorylation have provided the required tools to study these PTMs in HD models, identifying potential modulators of endogenous HTT phosphorylation (Cariulo et al., 2017) (Cariulo, Verani et al., 2019). Amongst such modulators are kinases capable of increasing phosphorylation at HTT T3 or S13/S16 (Bowie, Maiuri et al., 2018, Bustamante, Ansaloni et al., 2015, Chiki, Ricci et al., 2021, Ochaba, Fote et al., 2019, Thompson et al., 2009) as well as pharmacological tools for proof-of-concept studies (Alpaugh, Galleguillos et al., 2017, Atwal, Desmond et al., 2011, Bowie et al., 2018, Cariulo et al., 2019). Efforts to identify kinases and/or phosphatases associated with HTT phosphorylation at T3 and S13/S16 have so far led to the identification of IKBKB (Bustamante et al., 2015, Ochaba et al., 2019, Thompson et al., 2009), TBK1 (Hegde, Chiki et al., 2020) and PP1 (Branco-Santos et al., 2017) as tool enzymes for T3 (IKBKB, PP1) and S13/S16 (IKBKB, TBK1) regulation. Of these, IKBKB and/or IKK signalling have been mechanistically associated with mutant HTT toxicity (Khoshnan, Ko et al., 2009, Khoshnan, Ko et al., 2004, Khoshnan & Patterson, 2011, Khoshnan, Sabbaugh et al., 2017, Thompson et al., 2009), *HTT* transcriptional regulation (Becanovic, Norremolle et al., 2015) and modulation of HTT S13 phosphorylation in vivo (Ochaba et al., 2019). This evidence, together with the known amenability to pharmacological modulation, makes IKK pathway kinases (and particularly IKBKB) strong tool candidates that can elucidate the mechanisms involved in regulating mutant HTT S13/S16 phosphorylation (Atwal et al., 2011, Ochaba et al., 2019).

Studies using phosphor-mimetic mutations or bona-fide HTT phosphorylation have however suggested that increased phosphorylation at T3 or S13/S16 would need to be achieved (Atwal, Xia et al., 2007, Cariulo et al., 2017, Chiki A., 2017, DeGuire S.M., 2017, DeGuire et al., 2018, Gu, Greiner et al., 2009, Zheng et al., 2013), thereby pointing at the need to develop ways to increase (rather than inhibit) the activity of relevant kinases. The paradoxical observations that IKBKB can directly phosphorylate HTT at S13/S16 (Thompson et al., 2009) and that IKBKB inhibitors can achieve the same effect (Atwal et al., 2011) suggest a level of complexity which requires further investigation, examining the role of the distinct IKK signalling pathways downstream of IKBKB in the regulation of HTT N17 phosphorylation. IKK signalling is amongst the better understood kinase signalling pathways, and includes signalling by the IKK complex (IKBKB homodimers or IKBKB/IKBKA heterodimers in complex with IKBKG) regulating the activity of the transcription factor NF-kb (Hacker & Karin, 2006, Israel, 2010, Perkins, 2007, Yu, Lin et al., 2020), as well as non-IKK complex mediated signalling pathways, typically mediated by the non-canonical IKK kinases IKBKE and TBK1, the best understood of which acts via the nuclear factors IRF3/IRF7 (Balka, Louis et al., 2020, Clark, Peggie et al., 2011, Fitzgerald, McWhirter et al., 2003, Hacker & Karin, 2006, Sharma, tenOever et al., 2003, Yum, Li et al., 2021). While published evidence implicates IKBKB/IKBKA and canonical (NF-kb) mediated signalling in contributing to mutant HTT-mediated pathology and in *HTT* transcriptional regulation (Becanovic et al., 2015, Khoshnan & Patterson, 2011), little is known of the signalling downstream of IKBKB leading to HTT N17 phosphorylation.

Another level of complexity is provided by the presence of phosphorylation at T3 and S13/S16, which appear to functionally cross-talk at least in vitro (DeGuire et al., 2018). As IKBKB has been described capable of phosphorylating both these HTT phosphor-epitopes (Bustamante et al., 2015, Thompson et al., 2009), it is important to investigate the effects of T3 and S13/S16 phosphorylation by IKBKB from the perspective of cross-talk, exploring how T3 modification affects S13/S16 phosphorylation and vice-versa.

Here, we have addressed these open questions using novel ultrasensitive assays for measuring endogenous levels of S13 HTT phosphorylation (pS13 HTT). We confirm, for the first time on endogenous HTT, that IKBKB can quantitatively increase pS13 HTT levels in human cells and does so in a manner dependent on its catalytic activity, as determined by mutations inactivating its kinase domain and selective pharmacological inhibitors. We further uncover a role for phosphatases and in particular for PP2A in regulating the ability of IKBKB to modulate pS13 HTT levels. Interestingly, we found that the ability of IKBKB to increase pS13 HTT levels in human cells is shared with IKBKE but not with the other canonical IKK kinase, IKBKA, and used IKBKB mutants to define that monomeric as well as Nemo binding-incompetent IKBKB remain capable of increasing pS13 HTT levels, thus excluding a role for the IKK complex and canonical IKK signalling. Coherently, pS13 HTT modulation by IKBKB was found to be associated with activation of a non-canonical IKK pathway involving IRF3 activation rather than through IKBA (canonical IKK signalling) activation. Functionally, the modulation of pS13 HTT levels by IKBKB is associated with a robust decrease of aggregation in a commonly employed surrogate cell model of mutant HTT aggregation. Finally, we explored the role of residue cross-talk, providing the first evidence that T3 mutation affects basal and IKBKB induced pS13 HTT levels, an effect which translates functionally also in mutant HTT aggregation in cells. The identification of non-canonical IKK signalling as a regulator of HTT N17 phosphorylation and the finding that T3 and S13 phosphorylation may be subject to cross-talk represent major advances towards the understanding of approaches aiming at reducing mutant HTT pathology through increased N17 phosphorylation, and provide novel insights towards the mechanistic understanding of N-terminal phosphorylation, regulation and downstream pathophysiological implications.

## Results

### IKBKB modulation of endogenous pS13 HTT levels is regulated by okadaic acid-sensitive phosphatases

IKBKB can phosphorylate HTT N-terminal protein fragments in vitro (Thompson et al., 2009) and its overexpression can increase pS13 HTT levels in cells expressing HTT N-terminal fragments (Bustamante et al., 2015, Thompson et al., 2009). More recently, endogenous IKBKB was shown capable of regulating pS13 HTT levels in mice as determined by semiquantitative methods (Ochaba et al., 2019). Paradoxically, other studies showed that pharmacological inhibition of IKBKB leads to the same results in cells (Atwal et al., 2011). We sought to investigate this discrepancy by interrogating the role of phosphatases in regulating the effects of IKBKB on pS13 HTT levels. Phosphatases may affect pS13 HTT levels in different ways, including direct dephosphorylation of pS13 as well as regulation of the catalytic activity, localization and stability of S13 kinases and phosphatases. Of the different pharmacological tools available to probe phosphatase activity, okadaic acid (OA), a somewhat selective inhibitor of PP1 and PP2A (Dounay & Forsyth, 2002), has been previously used to demonstrate an increase in nuclear accumulation of N-terminal HTT fragments in cells associated with increased S16 HTT phosphorylation (Havel, Wang et al., 2011). Interestingly, both PP1 and PP2A have been associated with a role in regulating N-terminal HTT aggregation and phosphorylation (Branco-Santos et al., 2017, Metzler, Gan et al., 2010), while PP2A is a known regulator of IKBKB catalytic activity and downstream signalling (DiDonato, Hayakawa et al., 1997). We therefore used OA to investigate the role of PP1/PP2A in regulating basal and IKBKB-induced pS13 HTT levels. For these mechanistic studies, we chose the human cell line HEK293T, being widely employed as a reductionistic model to study HTT PTMs and their biology (Bustamante et al., 2015, Cong, Held et al., 2011, Daldin, Fodale et al., 2017, Fodale, Kegulian et al., 2014, Huang, Lucas et al., 2015, Schilling, Gafni et al., 2006, Zheng et al., 2013).This cell line expresses HTT protein endogenously at significant levels (Cariulo et al., 2017, Ratovitski, O’Meally et al., 2017) and the HTT protein is phosphorylated at T3 and S13/S16 under both endogenous and overexpression conditions (Cariulo et al., 2017, Cariulo et al., 2019), indicating the presence of the relevant signalling pathways required to phosphorylate HTT at these residues. The ability of overexpressed IKBKB or an IKBKB carrying a catalytically inactivating mutation K44M (Mercurio, Zhu et al., 1997) to increase pS13 HTT levels in HEK293T cells overexpressing an N571 mutant HTT fragment was then assessed by Western blotting and the Singulex immunoassay (SMC assay) (Cariulo et al., 2019) in the presence or absence of OA at relevant concentrations (Havel et al., 2011) (Fig. 1, A and B respectively). We assessed pS13 HTT levels by Western immunoblotting using the previously characterized rabbit polyclonal antibody, specific for pS13 HTT (Cariulo et al., 2019). As shown in Fig. 1A, all HEK293T samples expressed N571 Q55 HTT protein at comparable levels. However, in the absence of OA little if any increase in pS13 HTT levels was observed in cells transfected with IKBKB as compared to cells transfected with the empty expression plasmid, in spite of robust expression of IKBKB. However, this IKBKB protein was slightly inactive as judged by the level of phosphorylation at residues S177/S181, which are auto-phosphorylation sites commonly employed as a readout of IKBKB activity (Israel, 2010). We reasoned that the absence of robust pS13 HTT modulation by IKBKB was a result of low IKBKB activity, potentially due to the activity of endogenous phosphatases in HEK293T cells. In the presence of OA, however, a robust increase in pS13 HTT levels was observed in samples transfected with IKBKB but not in those transfected with an IKBKB kinase dead construct (Fig. 1A), paralleled with a strong increase in pS177/pS181 IKBKB levels, indicative of robust kinase activity. Interestingly, basal levels of pS13 HTT, in the absence of transfected IKBKB, were not affected by OA treatment, suggesting that control of pS13 HTT levels by OA-sensitive endogenous phosphatases is not the predominant regulatory mechanism. When samples were analysed by quantitative SMC assays, the capacity of OA to reveal a robust (ca. 10-fold), catalytic activity-dependent induction of pS13 HTT levels by IKBKB was confirmed (Fig. 1B). We concluded that while basal levels of pS13 HTT in cells expressing N571 Q55 HTT fragments are not significantly altered by OA-mediated phosphatase inhibition, the capacity of IKBKB to increase pS13 HTT levels is exquisitely controlled by OA-sensitive phosphatases, due to regulation of IKBKB activity, as evidenced by IKBKB pS177/pS181 levels. We next sought to determine if we could confirm the catalytic activity-dependence of IKBKB in regulating pS13 HTT levels and the effect of OA using pharmacological rather than genetic means, on either overexpressed N571 Q55 HTT (Fig. 1, C and D) or on endogenously expressed HTT in HEK293T cells (Fig. 1, E and F). As shown in Fig. 1C, N571 Q55 pS13 HTT levels were again not significantly modulated by IKBKB overexpression in the absence of OA, and addition of Bay-65-1942, a selective IKBKB inhibitor (Atwal et al., 2011), was similarly unable to significantly modulate these levels. In the presence of OA, the capacity of IKBKB to increase pS13 HTT levels was revealed and, importantly, it was sensitive to selective pharmacological inhibition by Bay-65-1942 in a concentration-dependent manner (Fig. 1D). Substantially comparable results were obtained when the response of pS13 HTT levels expressed from the endogenous *HTT* locus to IKBKB and OA was investigated under the same conditions (Fig. 1, E and F), and the more quantitative SMC assay analysis on these samples confirmed the findings obtained by Western blotting (no OA: Fig. 1G; OA treatment: Fig. 1H). Therefore, we concluded that pS13 HTT levels, whether from the endogenous *HTT* locus or from an overexpressed HTT N-terminal fragment, could be robustly increased in response to IKBKB in a catalytic activity-dependent and phosphatase-sensitive manner. As OA is described as an inhibitor of PP1 and PP2A (Dounay & Forsyth, 2002) but exhibits greater selectivity towards the latter (Swingle, Ni et al., 2007), the role of PP2A on OA’s ability to regulate IKBKB activity (S177/S181 auto-phosphorylation) and its effects on pS13 HTT levels was further evaluated using a specific siRNA to silence endogenous PP2A in HEK293T cells. As shown in Fig. 2A, the PP2A silencing produced an increase in IKBKB auto-phosphorylation, indicative of increased IKBKB activity, concomitantly with increased pS13 HTT levels. The magnitude of the effects of PP2A silencing on pS13 HTT modulation by IKBKB was not comparable to that attained with OA, likely due to incomplete PP2A silencing obtained with RNAi and the difficulty of phenocopying pharmacological inhibition of a catalytic enzyme with RNAi (Weiss, Taylor et al., 2007). Coherent with the observations made with OA, basal pS13 HTT levels (absence of IKBKB overexpression) were not affected by PP2A RNAi. Quantitative analysis of pS13 HTT levels of overexpressed N571 HTT bearing 55 CAG repeats (Q55) by SMC assay essentially confirmed the results obtained by Western blotting (Fig. 2B). Consistent with a role for PP2A in regulating IKBKB activity, we were able to detect an interaction between overexpressed IKBKB and endogenously expressed PP2A in HEK293T cells by both immunoprecipitation (Fig. 2C) and an ELISA interaction assay (Fig. 2D). We next sought to determine if overexpression of PPP2CA, the catalytic subunit of PP2A, could affect the ability of IKBKB to influence pS13 HTT levels, an experiment which needed to be performed in the absence of OA and therefore against the background of endogenous inhibitory effects of PP2A on IKBKB. Using overexpression of N571 Q55 HTT and shorter transfection times (24 hrs) than those employed in Fig. 1, we were able to detect a reproducible, modest induction of pS13 HTT levels upon co-transfection with IKBKB, concomitant with significant S177/S181 IKBKB phosphorylation as a measure of IKBKB catalytic activity (Fig. 2E and F). Under these conditions, co-transfection of PPP2CA effectively abolished IKBKB induced phosphorylation at S177/S181 and pS13 HTT levels, consistent with the observations made using RNA knock-down. Collectively, the data suggest that the capacity of IKBKB to influence pS13 HTT levels in HEK293T cells is dependent on its catalytic activity, that the catalytic activity of IKBKB is regulated by OA-sensitive phosphatases and that PP2A contributes to IKBKB inducible, but not basal, pS13 HTT levels. These findings are fully consistent with the known role of PP2A in regulating IKBKB activity (Barisic, Strozyk et al., 2008, Tsuchiya, Osaki et al., 2017).

**Figure 1.**
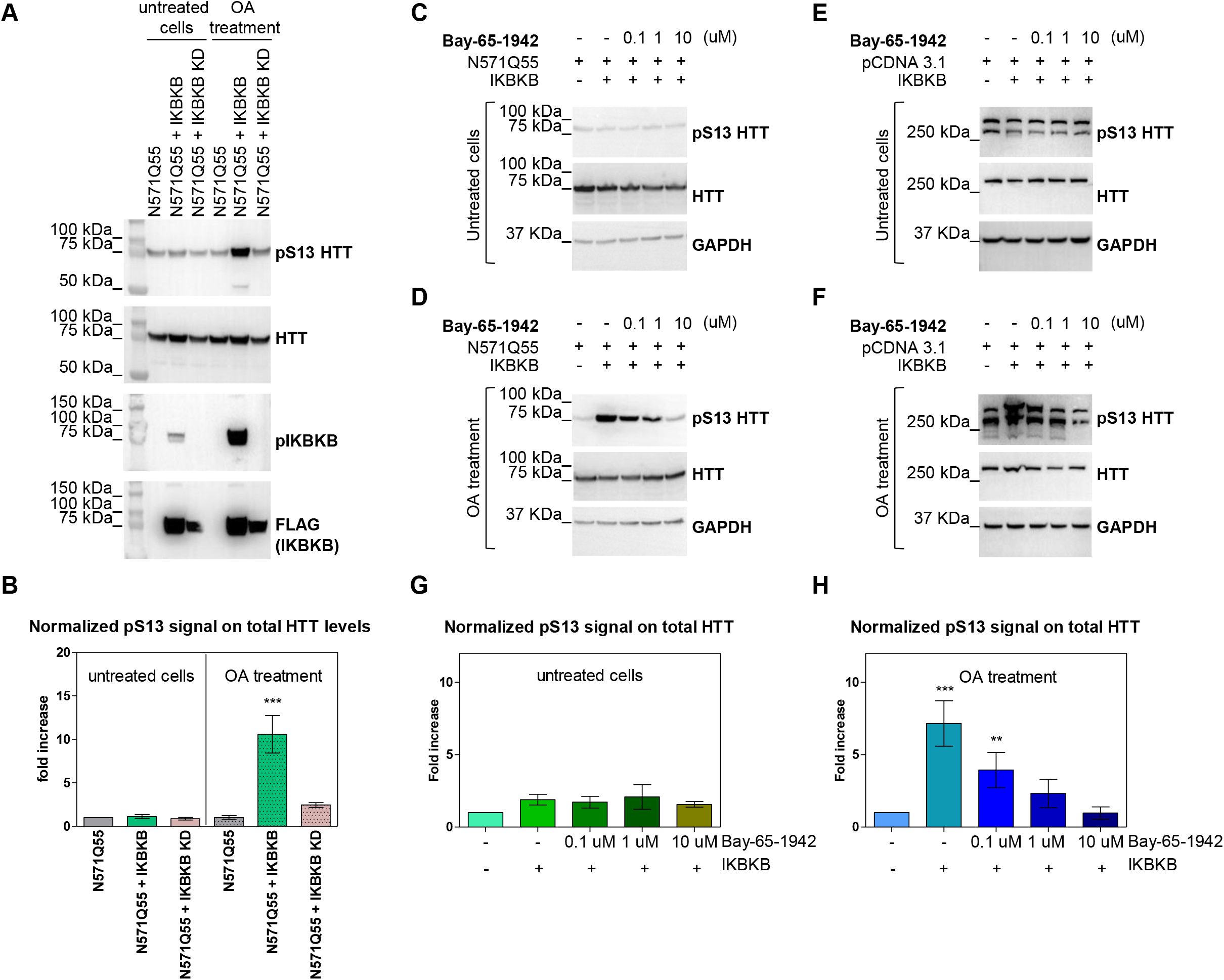
pS13 HTT levels are regulated by IKBKB and okadaic acid-sensitive phosphatases. A-B Treatment with okadaic acid (OA) of HEK293T cells overexpressing N571 Q55 HTT, with or without IKBKB or its kinase dead mutant IKBKB KD. (A) Western blotting showing robustly increased pS13 HTT levels as well as increased auto-phosphorylation of IKBKB at residues S177/S181 upon IKBKB but not IKBKB KD overexpression in combination with OA treatment. Kinase expression and HTT levels were assessed using anti-FLAG and mAb 2166 antibodies respectively. (B) Normalized pS13 signal on total HTT levels measured by SMC assay performed on the same lysates as in (A) confirming the Western blotting results. (Means and SDs were calculated on 3 biological replica. One-way analysis of variance, Dunnett’s Test (***P < 0.0001)). C-D Western blotting of HEK293T cells overexpressing N571 Q55 HTT with or without IKBKB treated with IKBKB-inhibiting-compound BAY-65-1942 at 0.1, 1 and 10 uM, without OA (C) and in combination with OA (D), demonstrating the capacity of IKBKB to increase pS13 HTT levels only in presence of OA, and of BAY-65-1942 compound to inhibit IKBKB’s activity on pS13 in a dose dependent way. Protein loading and HTT levels were assessed using anti-GAPDH and mAb 2166 antibodies respectively. E-F Same experimental conditions as in (C) and (D) evaluated on endogenous HTT in HEK293T cells confirming the overexpression results without OA (E) or in presence of OA (F). Protein loading and HTT levels were assessed using anti-GAPDH and mAb 2166 antibodies respectively. G-H SMC assay on same lysates as in (C)-(F) (means and SDs of n=2 overexpressed HTT merged with n=2 endogenous HTT experiments) without OA (G) and in combination with OA (H), confirming Western blotting results. (Means and SDs were calculated on 4 biological replica. One-way analysis of variance, Dunnett’s Test (**P < 0.001; ***P< 0.0001)).

**Figure 2.**
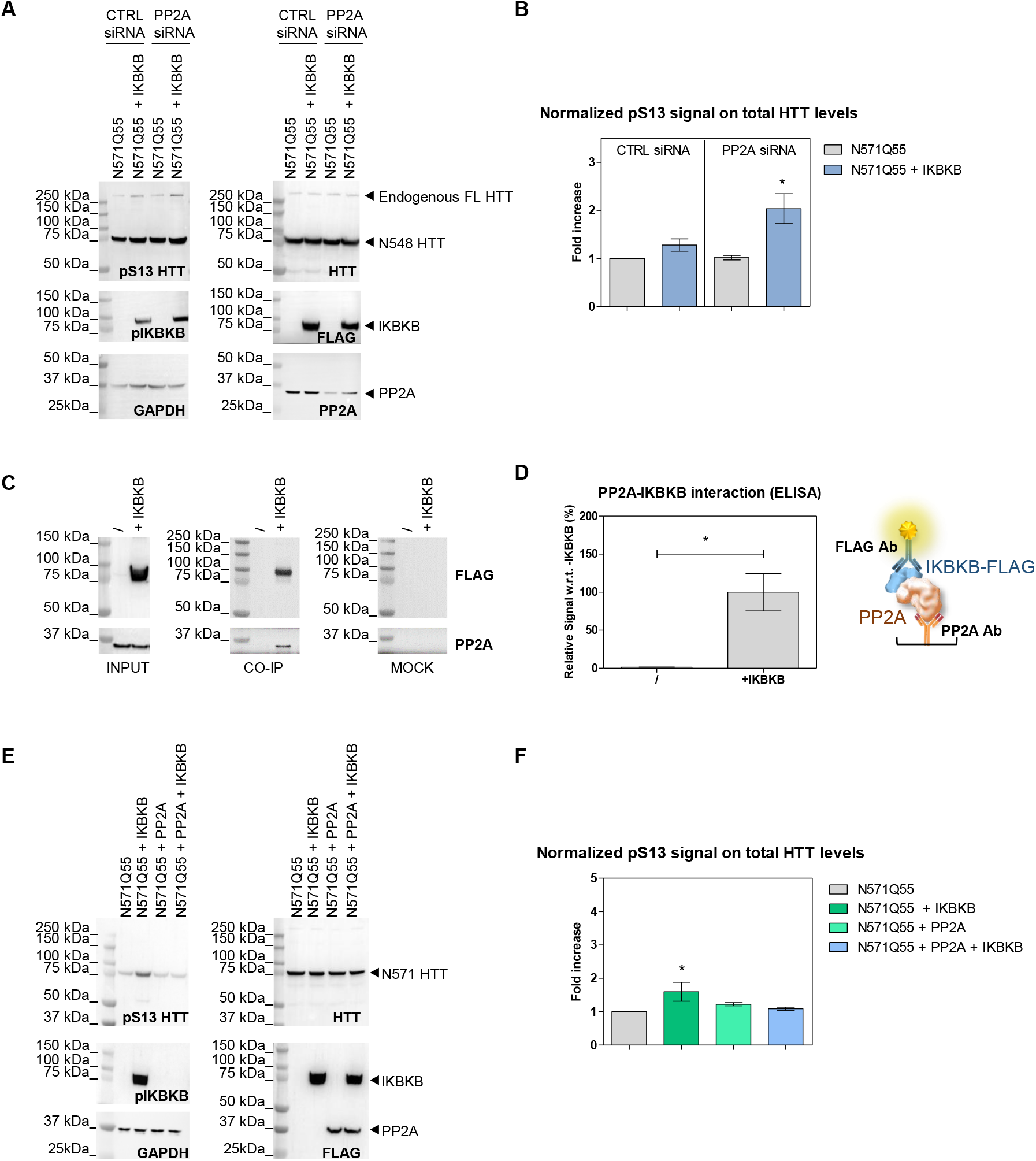
PP2A interferes on IKBKB’s capacity to phosphorylate HTT S13. A-B Endogenous PP2A knockdown consequent to specific siRNA treatment in HEK293T cells transiently co-transfected with N571 Q55 HTT and IKBKB produced an increased response to IKBKB compared to siRNA control. (A) Western blotting showing that IKBKB overexpression leads to higher phosphorylation on HTT S13 when PP2A is silenced (as shown by anti-PP2A Ab). Increased autophosphorylation of IKBKB on S177/S181 as a result of endogenous PP2A knockdown. Protein loading, kinase and phosphatase expression and HTT levels were assessed using anti-GAPDH, antiFLAG, anti-PP2A and mAb 2166 antibodies respectively. (B) Normalized pS13 signal on total HTT levels measured by SMC assay performed on the same lysates as in (A) confirming Western blotting results. (Means and SDs were calculated on 3 biological replica. One-way analysis of variance, Dunnett’s Test (*P < 0.05)). C-D Endogenous PP2A in HEK293T cells interacts with IKBKB. (C) Western blotting of HEK293T cells lysates overexpressing IKBKB FLAG-tagged (INPUT) pulled-down by anti-FLAG antibody (CO-IP) or unrelated anti-GFAP antibody (MOCK), and revealed by anti-FLAG and anti-PP2A antibodies. Anti-FLAG and anti-PP2A signals in CO-IP at the expected molecular weights demonstrate the PP2A-IKBKB interaction. (D) ELISA assay using anti-PP2A as capture antibody and anti-FLAG as detection antibody performed on HEK293T cell lysates transfected with/without IKBKB FLAG-tagged confirms the interaction between endogenous PP2A and IKBKB. (Means and SDs were calculated on 3 biological replica. T-test (*P < 0.05)). E-F IKBKB’s capacity to influence pS13 HTT levels is affected by PP2A. (E) Western blotting of HEK293T cells overexpressing N571 Q55 HTT with or without IKBKB FLAG-tagged, PP2A FLAG-tagged or the combination of both. The lack of up-regulation of pS13 HTT levels by IKBKB when PP2A is overexpressed demonstrates that PP2A can regulate IKBKB’s activity (as confirmed by the absence of IKBKB-auto-phosphorylation in the same condition) rather than dephosphorylating HTT S13 (as shown by lack of modulation of pS13 HTT levels in the presence of PP2A). Protein loading, kinase and phosphatase expression and HTT levels were assessed using anti-GAPDH, anti-FLAG and mAb 2166 antibodies respectively. (F) Normalized pS13 signal on total HTT levels measured by SMC assay, performed on the same lysates as in (E) confirming the Western blotting results. (Means and SDs were calculated on 3 biological replica. One-way analysis of variance, Dunnett’s Test (*P < 0.05)).

### IKBKB modulation of endogenous pS13 HTT levels involves a non-canonical IKK pathway associated with IRF3 signalling

IKK kinases have been extensively studied owing to their involvement in multiple areas of human disease, including inflammation, immunity and cancer (Clark et al., 2011, Hacker & Karin, 2006, Israel, 2010, Mercurio et al., 1997, Perkins, 2007, Sharma et al., 2003, Zandi, Rothwarf et al., 1997). Several functional mutants have been extensively characterized to probe the involvement of IKBKB in IKK pathways, which broadly can be grouped into IKK complex-mediated (canonical) and non-canonical. Canonical IKK signalling involves the formation of a functional complex involving IKBKB, IKBKA and IKBKG/Nemo, and results in IKBA phosphorylation and degradation, release of NF-kb from its inhibitor IKBA, and translocation of NF-kb to the nucleus. NF-kb executes its role as a transcription factor (Israel, 2010, Yu et al., 2020), regulating the expression of a number of genes involved in, amongst others, inflammation but also HTT itself (Becanovic et al., 2015). In contrast, non-canonical IKK signalling is largely IKBKG independent and includes pathways of which the best understood are those triggered by the non-canonical IKK kinases, TBK1 and IKBKE, which involve IRF factors (IRF3/7) rather than NF-kb as transcriptional effectors (Balka et al., 2020, Fitzgerald et al., 2003, Hacker & Karin, 2006, Yum et al., 2021). A range of IKBKB mutants have been generated and characterized which can be leveraged to genetically investigate the mechanism of IKBKB regulation of pS13 HTT levels. The HEK293T cell model, with its simplicity and amenability to manipulation, and the observation that OA unmasks the ability of IKBKB to modulate pS13 HTT by inhibiting endogenous phosphatases including PP2A, provides an appropriate environment for these mechanistic investigations. Further validation and hypothesis building, with regard to relevance for HD needs to be subsequently tested in other, more complex contexts. Equipped with IKBKB and a set of IKBKB mutants (Fig. 3A) including a catalytically inactive (IKBKB KD) variant (Mercurio et al., 1997), an IKBKG/Nemo binding-incompetent (IKBKB NBD) mutant (May, Marienfeld et al., 2002), a variant with decreased kinase activity through phosphor-mimetic mutations at residues S177/S181 (IKBKB S177E/S181E) (Huynh, Boddupalli et al., 2000, Kishore, Huynh et al., 2002, Liu, Misquitta et al., 2013, Mercurio et al., 1997, Polley, Huang et al., 2013) and an IKBKB mutant incapable of dimerization (IKBKB LZ) (Hauenstein, Rogers et al., 2014), we set out to dissect the mechanisms through which IKBKB increases pS13 HTT levels in the HEK293T model. All mutants were C-terminally FLAG-tagged to facilitate comparative analysis of expression levels. We co-transfected IKBKB and the indicated mutants in HEK293T cells together with an expression plasmid encoding N571 Q55 HTT, in the presence or absence of OA, and determined their activity on pS13 HTT levels. As shown in Fig. 3B, in the absence of OA treatment, IKBKB did not produce a robust increase in pS13 HTT levels, as previously observed, as did none of the IKBKB mutants in spite of varying expression levels of these latter (see anti-FLAG blotting). Interestingly in the presence of OA, IKBKB and IKBKB mutant incapable of forming a complex with IKBKG/Nemo (IKBKB NBD) or the IKBKB mutant incapable of dimerization (IKBKB LZ) were still capable of increasing pS13 HTT levels, with a concomitant increase in S177/S181 phosphorylation, consistent with an IKK complex (IKBKG/IKBKA)-independent mechanism. This was evident despite the disparity in expression levels of the different constructs, which became evident with the kinase dead IKBKB (IKBKB KD) and the IKBKG/Nemo binding-incompetent IKBKB (IKBKB NBD). The latter construct still produced a substantial increase in pS13 HTT levels, in spite of significantly lower expression levels with respect to for instance IKBKB. These samples were also analysed using SMC assays, producing essentially comparable results to those obtained by Western blotting (Fig. 3C). The data indicated that, in HEK293T cells, IKBKB regulates pS13 HTT levels through a non-canonical, IKK complex-independent mechanism, dependent on the catalytic activity of IKBKB which itself is regulated by protein phosphatases including PP2A. We sought to confirm and further extend these findings by analyzing the effects of overexpression of IKBKB and its mutants on endogenous pS13 HTT levels in comparison with overexpression of a canonical (IKBKA) or a non-canonical (IKBKE) IKK kinase, and of the non-catalytic scaffolding component of the canonical IKK complex, IKBKG/Nemo. As shown in Fig. 4A, in the absence of OA overexpressed IKBKB, IKBKB NBD and IKBKB LZ were able to modestly increase endogenous pS13 HTT levels, while IKBKB KD and the IKBKB S/E mutant (decreased kinase activity) were not, as observed in cells expressing the N571 Q55 HTT protein (Fig. 3, B and C). Interestingly, overexpression of the non-canonical kinase IKBKE but not of the canonical kinase IKBKA or of IKBKG, the non-catalytic scaffolding component of the canonical IKK complex, was also able to increase endogenous pS13 HTT levels (Fig. 4A and B). In the presence of OA these effects were greatly enhanced, as expected (Fig. 4C and D). These results confirmed that non-canonical IKK signalling, but not signalling mediated by the canonical IKK complex, was responsible for regulating pS13 HTT levels. To further support this finding, we examined the phosphorylation of IRF3, a characterized effector of non-canonical IKK signalling, which upon phosphorylation by non-canonical IKK kinases is activated and acts as the transcriptional mediator of the pathway (Fitzgerald et al., 2003, Hacker & Karin, 2006). Indeed, we observed complete correlation between pS13 HTT phosphorylation and IRF3 phosphorylation (Fig. 4A and 4C). Collectively, the effects of IKBKB on pS13 HTT levels expressed from the endogenous *HTT* locus or from overexpressed HTT fragments are dependent on its catalytic activity, which is regulated by phosphatases including PP2A, can be phenocopied by non-canonical IKK kinases such as IKBKE but not by canonical IKK kinase IKBKA and, consistently, are mediated by a non-canonical IKK pathway associated with IRF3 activation.

**Figure 3.**
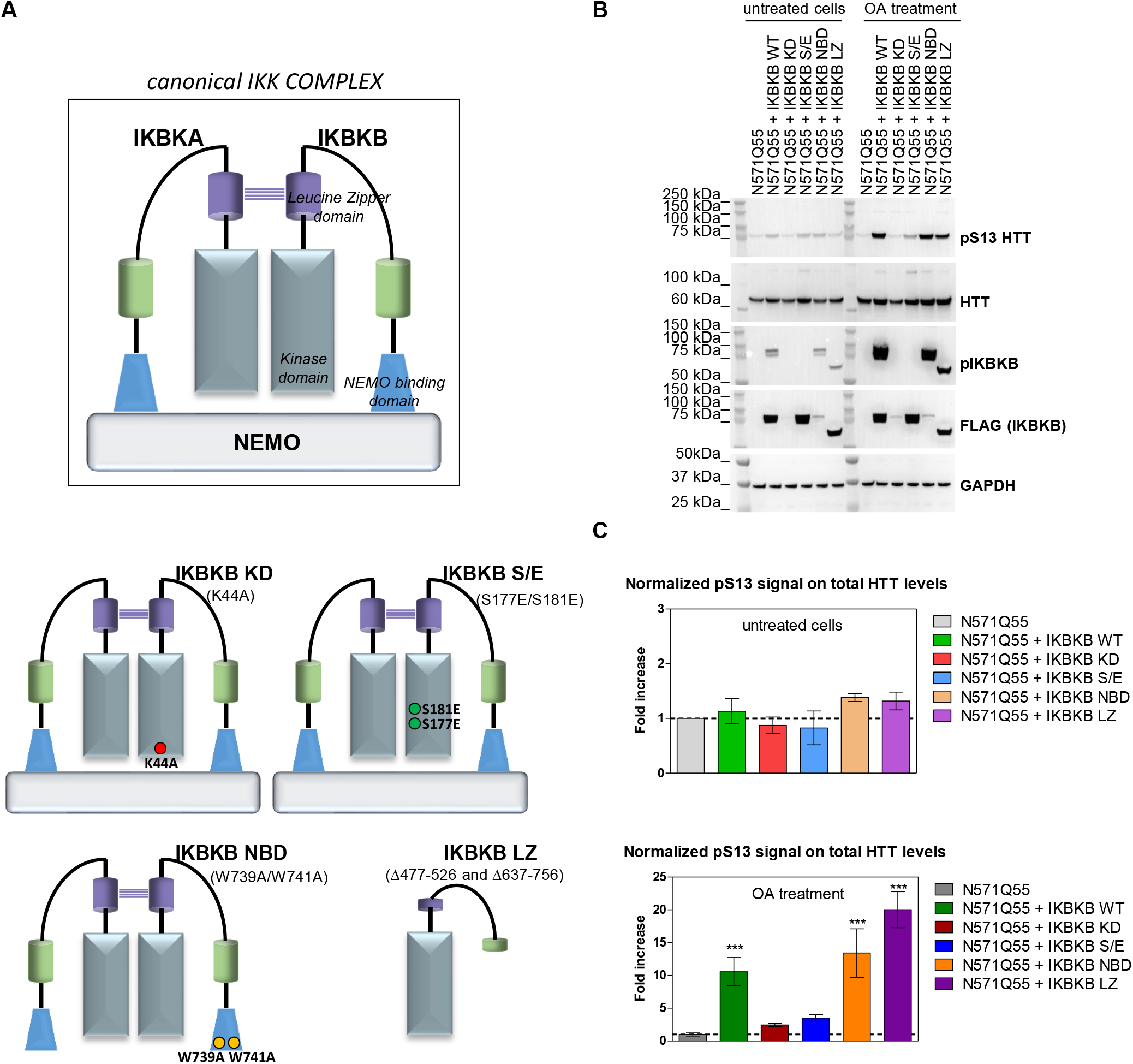
IKBKB effects on pS13 HTT levels are independent of IKBKB’s activity in the canonical IKK complex. A Schematic representation of IKBKB mutant constructs employed here. B-C (B) Western blotting of HEK293T cells overexpressing N571 Q55 HTT with or without IKBKB WT and IKBKB mutants, in presence or absence of OA treatment. pS13 HTT levels increased only in the presence of the IKBKB WT, the IKBKB NBD and the IKBKB LZ mutants when cells were treated with OA, indicating that the canonical IKK pathway is not involved in regulation of pS13 by IKBKB. Protein loading, kinases expression and activity, as well as HTT levels were assessed using anti-GAPDH, antiFLAG, anti-pIKBKB and mAb 2166 antibodies respectively. (C) Normalized pS13 signal on total HTT levels measured by SMC assay, performed on the same lysates as in (B) confirming the Western blotting results. (Means and SDs were calculated on 3 biological replica. One-way analysis of variance, Dunnett’s Test (***P < 0.0001)).

**Figure 4.**
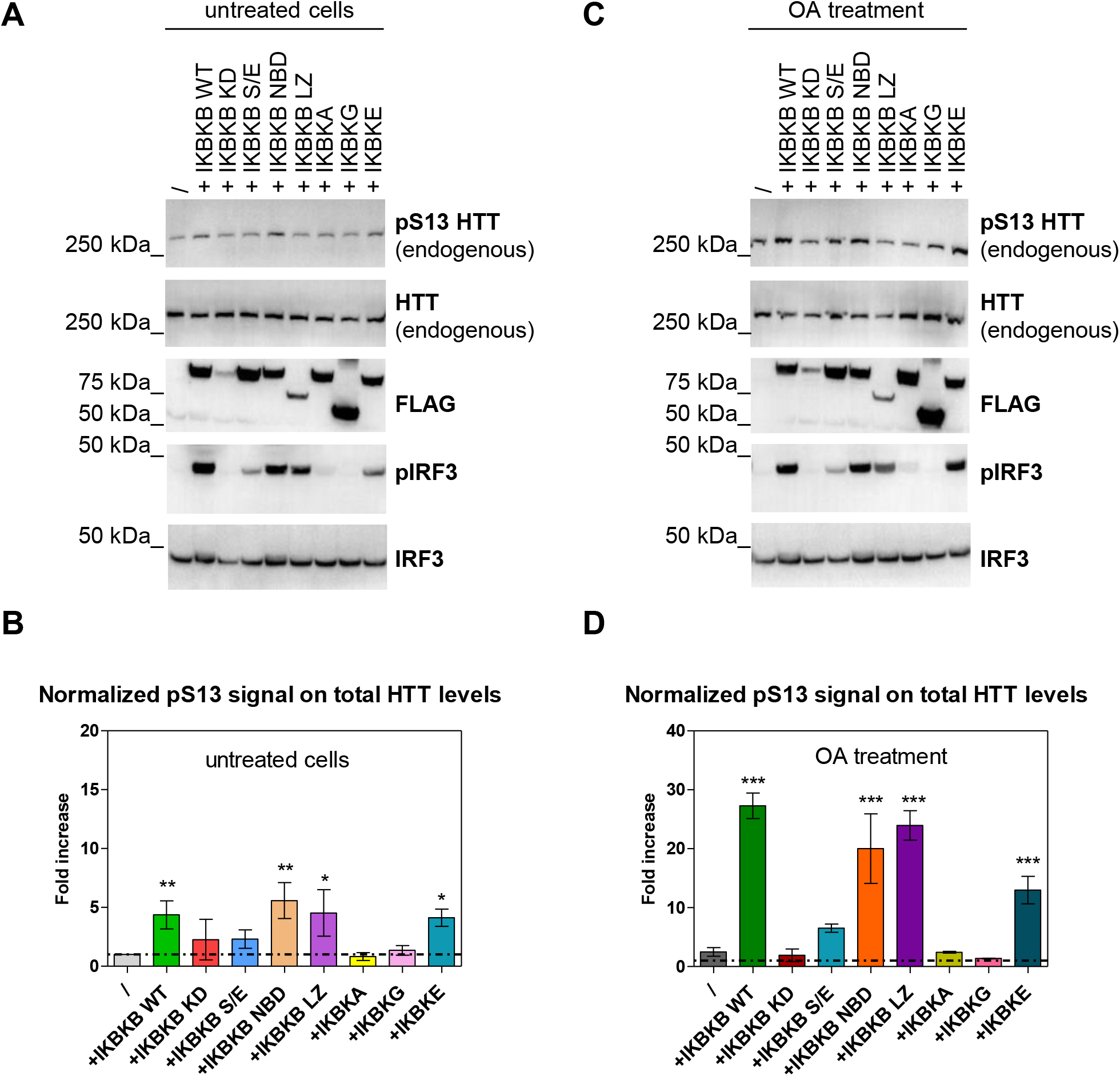
IKBKB regulation of endogenous pS13 HTT levels is associated with an IRF3 mediated, non-canonical IKK pathway. A-B HEK293T cells overexpressing IKBKB WT and IKBKB mutants, IKBKA, IKBKG and IKBKE in absence of OA. (A) Western blotting showing a modest increase of endogenous pS13 HTT levels upon expression of IKBKB WT, IKBKB NBD mutant, IKBKB LZ mutant as well as of the non-canonical IKBKE kinase. The non-canonical IRF3 pathway is involved in pS13 HTT regulation as demonstrated by the increase of endogenous phosphor-IRF3. Kinases expression, HTT levels and endogenous IRF3 expression were assessed using anti-FLAG, mAb 2166 and anti-IRF3 antibodies respectively. (B) Normalized pS13 signal on total HTT levels measured by SMC assay, performed on the same lysates as in (A) confirming the Western blotting results. (Means and SDs were calculated on 3 biological replica. One-way analysis of variance, Dunnett’s Test (*P < 0.05; **P < 0.001)). C-D HEK293T cells overexpressing IKBKB WT and IKBKB mutants, IKBKA, IKBKG and IKBKE in presence of OA. (C) Western blotting showing that the presence of OA greatly enhances the effects described in (A) and (B), coherent with effects on pS13 HTT being dependent on the catalytic activity of IKBKB. (D) Normalized pS13 signal on total HTT levels measured by SMC assay, performed on the same lysates as in (C) confirming the Western blotting results. (Means and SDs were calculated on 3 biological replica. One-way analysis of variance, Dunnett’s Test (***P < 0.0001)).

IKBKB is known to directly phosphorylate purified, isolated HTT exon 1 protein *in vitro* (Thompson et al., 2009) but no evidence of a direct interaction between IKBKB and HTT has been demonstrated in cellular or animal models. We therefore asked whether the kinase exerted its activity on S13 in a direct way, through interaction with HTT in a cellular context, by applying a quantitative protein-protein interaction assays (TR-FRET and ELISA) to HEK293T cells co-expressing N571 Q55 HTT construct with or without S13A/S16A phosphor-abrogative mutations and IKBKB. As shown in Fig. 5 (A, B and C), IKBKB was able to interact with HTT N571 Q55 protein and this interaction was independent of the phosphorylation status of S13, as the abrogation of S13 phosphorylation (S13A/S16A mutant) did not impact a direct interaction. On the other hand the interaction was dependent on the catalytic activity of IKBKB, as the catalytically inactive mutant (KD) interacted with the HTT protein to a lesser extent (Fig. 5D). Based on these data we conclude that IKBKB can interact with HTT in this cell context, and that this interaction is dependent on the integrity of the catalytic domain of IKBKB but independent of IKBKB’s HTT substrate phosphorylation.

**Figure 5.**
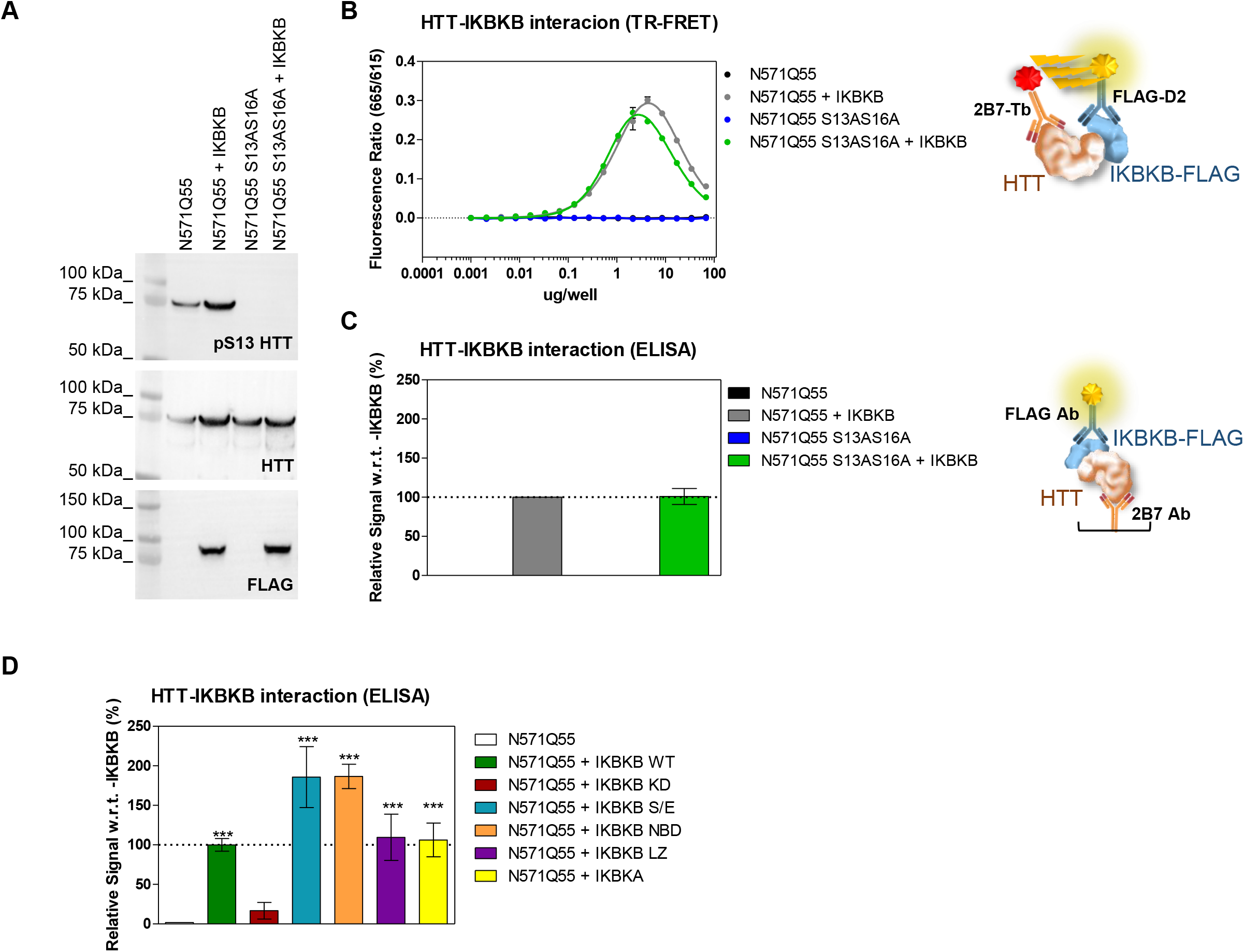
IKBKB interacts with HTT in cells in a manner dependent on its catalytic activity but independent of HTT phosphorylation at S13. A-C (A) Western blotting of HEK293T cells overexpressing N571 Q55 HTT WT or S13A/S16A phosphor-abrogative mutant with or without IKBKB FLAG-tagged. pS13 HTT levels, IKBKB expression and HTT levels were assessed using anti-pS13, anti-FLAG and mAb 2166 antibodies respectively. (B) TR-FRET protein-protein interaction assay employing 2B7-Tb/anti-FLAG-D2 antibody pair showing a strong interaction of N571 Q55 HTT WT with IKBKB but also with the phosphor-abrogative N571 Q55 HTT S13A/S16A mutant. (C) ELISA protein-protein interaction assay employing 2B7 as capture and anti-FLAG as detection antibodies used as orthogonal readout confirming the data in (B). D ELISA protein-protein interaction assay of HEK293T cells overexpressing N571 Q55 HTT WT and IKBKB WT or IKBKB KD, IKBKB S/E, IKBKB NBD, IKBKB LZ or IKBKA, showing that the interaction of N571 Q55 HTT with IKBKB is robustly affected only in the presence of the kinase dead mutation and therefore this interaction is dependent on the integrity of IKBKB’s catalytic domain. (Means and SDs were calculated on 3 biological replica. One-way analysis of variance, Dunnett’s Test (***P < 0.0001)).

### IKBKB influences mutant HTT aggregation through increased S13 HTT phosphorylation and cross-talk between T3 and S13

One of the key functional consequences of polyQ expansion believed to contribute to HD pathology is misfolding and aggregation of short (exon 1 encoded) N-terminal HTT fragments, produced either from misplicing and/or proteolysis (Bates, 2003, Bates et al., 2015, Franich, Hickey et al., 2019, Gipson, Neueder et al., 2013, Landles et al., 2010, Neueder, Landles et al., 2017, Sathasivam et al., 2013, Saudou & Humbert, 2016, Weiss, Trager et al., 2012). Although mutant HTT aggregation is invariably (but to different extents) detectable in HD mouse models (Menalled & Brunner, 2014, William Yang & Gray, 2011), this process is not easily reproduced in cellular models of full length mutant HTT. To address this issue, expression of the exon 1 fragment of mutant HTT in cells, often tagged with fluorescent proteins, is generally employed as a surrogate mutant HTT aggregation model (Atwal et al., 2007, Bates et al., 2015, Branco-Santos et al., 2017, Cooper, Schilling et al., 1998, Li, Liu et al., 2016, Li & Li, 1998, Olshina, Angley et al., 2010, Ross & Tabrizi, 2011, Zheng et al., 2013). Indeed, mutant HTT aggregation and toxicity in cellular and animal models of HD is robustly affected by phosphor-mimetic mutations at S13/S16 as well as by other mutations within the N17 domain (Atwal et al., 2007, Branco-Santos et al., 2017, Greiner & Yang, 2011, Gu et al., 2009, Zheng et al., 2013). Although some pharmacological tools demonstrated to increase pS13/pS16 HTT levels have shown benefit in animal and cellular models of HD, modifying different phenotypes including mutant HTT aggregation (Alpaugh et al., 2017, Bowie et al., 2018, Di Pardo, Maglione et al., 2012), the lack of suitable direct genetic modulators of pS13/pS16 HTT levels have so far limited the capacity to determine if effects of phosphor-mimetic mutations are indeed a true phenocopy of increased phosphorylation. Together with the data presented here, the identification of IKBKB as a kinase capable of increasing S13/S16 HTT phosphorylation (Thompson et al., 2009) and the evidence supporting its role as a physiologically relevant modulator of N-terminal HTT phosphorylation (Ochaba et al., 2019) points to IKBKB as a suitable tool to probe the role of increased phosphorylation on mutant HTT aggregation. In order to do this, we first implemented a surrogate cellular model of mutant HTT aggregation, based on overexpression of HTT exon 1 bearing an expanded polyQ (Q72) repeat, C-terminally fused to GFP, in HEK293T cells (Fodale et al., 2014). To validate the model, we sought to confirm the previously reported effects of N17 mutations on mutant Exon 1 HTT (mut Ex1 HTT) aggregation, where phosphor-mimetic mutations at S13/S16 resulted in robustly decreased aggregation (Atwal et al., 2007, Branco-Santos et al., 2017, Greiner & Yang, 2011), using previously validated and characterized constructs (Cariulo et al., 2017, Fodale, Boggio et al., 2017). As illustrated in Fig. 6A and B, consistent with published data (Branco-Santos et al., 2017, Gu et al., 2009) S13A/S16A and S13D/S16D double mutations produced dramatic and opposite effects on mut Ex1 HTT aggregation (negative controls are shown in Fig. S1). These data are in agreement with the proposed role for phosphomimesis in decreasing mut Ex1 HTT aggregation (Greiner & Yang, 2011). Conversely, T3A and T3D mutations also decreased aggregation (Fig. 6), but to the same extent indicative that the effect cannot be reconciled with phosphomimesis, contrary to what observed for S13/S16. In agreement with the results obtained using the independent T3D and S13D/S16D mutations, the triple T3D/S13D/S16D mutant produced a robust decrease in aggregates. Interestingly, a triple T3A/S13A/S16A mutation resulted in increased aggregation, comparable to that attained by the double S13A/S16A mutation (Fig. 6), suggesting that the beneficial effect of the T3A mutation on aggregation requires the presence of intact S13/S16 residues. One possibility is that mutation of T3 results in increased phosphorylation of S13/S16, and that the decreased aggregation observed with T3 mutation at least in part is mediated by increased pS13/pS16 levels. To investigate this aspect, we measured basal and IKBKB-induced pS13 HTT levels by densitometry analysis of Western blotting in cells transfected with HTT Ex1 Q72-EGFP and with T3A or S13A/S16A mutations (Fig. 7, A and B). As previously observed, a modest effect of IKBKB co-transfection on pS13 HTT levels was observed in cells transfected with the HTT Ex1 Q72-EGFP construct, owing to the absence of OA treatment. Indeed, the presence of a T3A mutation significantly increased basal pS13 HTT levels, by about 5-fold and correspondingly potentiated the effects of IKBKB on pS13 HTT levels relatively to the protein expressed from the unmodified construct (Fig. 7, A and B). We also measured pT3 HTT levels to determine if S13/S16 modification would affect pT3 HTT levels, which was not the case (Fig. 7, A and C), however confirming the ability of IKBKB to increase pT3 HTT levels (Bustamante et al., 2015). Therefore, in this reductionistic, surrogate model of mutant HTT aggregation we were able to confirm previous reports of the capacity of phosphor-mimetic S13/S16 mutations to reduce aggregation in cells, pointing to the role for T3 in regulating basal- and IKBKB-induced pS13 HTT levels. This also provides an explanation for the observed effects of T3A mutations to reduce aggregation in an S13/S16 dependent manner (Fig. 6). We next proceeded with examining the role of IKBKB-regulated pS13 HTT levels in the aggregation process. As previously shown, IKBKA and IKBKB S/E co-expression did not modulate pS13 HTT levels and as reported in Fig. 8A and B, also failed to modulate aggregation while IKBKB robustly reduced HTT Ex1 Q72-EGFP aggregation in a catalytic dependent manner, demonstrated by lack of effects of IKBKB KD. As this effect could be mediated directly through N17-phosphorylation dependent or independent mechanisms, we interrogated the relative importance of S13/S16 and T3 for the IKBKB mediated reduction in mutant HTT aggregation using the available mutations. As illustrated in Fig. 8C and D, the effects of IKBKB on HTT Ex1 Q72-EGFP aggregation were exquisitely dependent on the presence of intact S13/S16 residues, with T3 mutation further enhancing the capacity of IKBKB to influence the aggregation process, although T3A HTT Ex1 Q72-EGFP showed an already reduced propensity to aggregate (Fig. 6). We conclude that, in this surrogate cellular model of mutant HTT aggregation, IKBKB can reduce a pathologically relevant phenotype in a manner dependent on its catalytic activity and on the presence of intact S13/S16 residues, which directly links the ability of IKBKB to regulate pS13 HTT levels with its capacity to reduce aggregation. To our knowledge this is the first demonstration that increasing S13 phosphorylation phenocopies the effects of S13/S16 phosphor-mimetic mutations.

**Figure 6.**
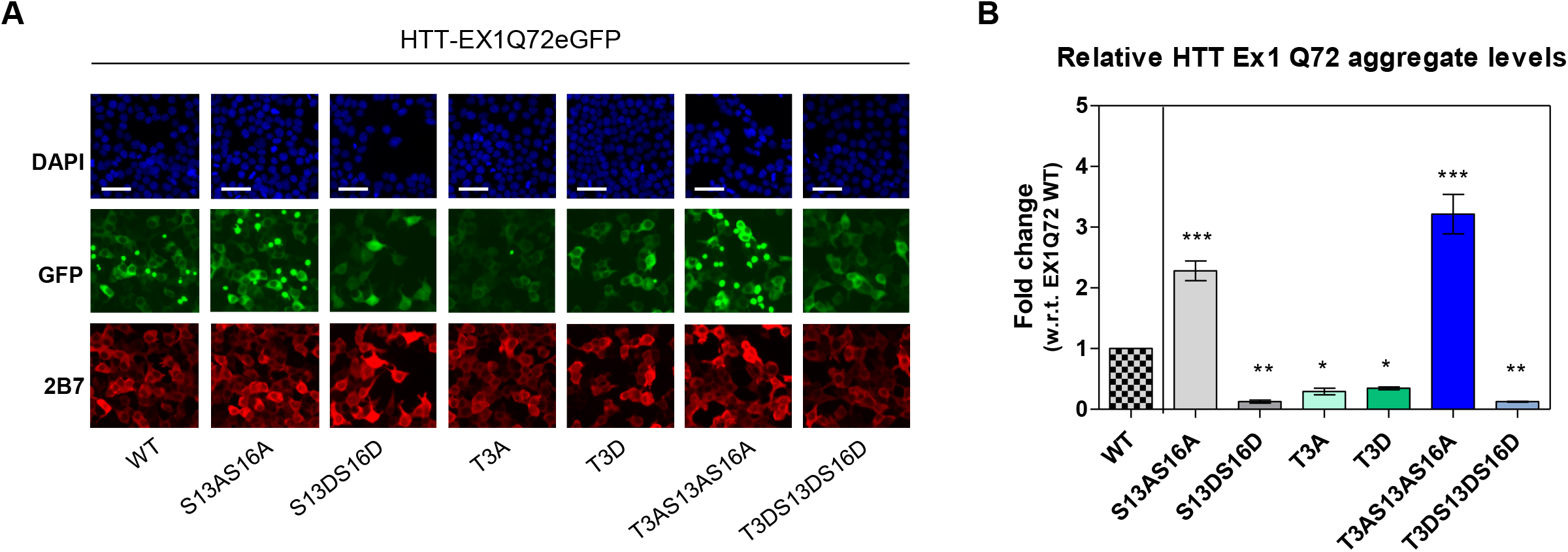
HTT N-terminal phosphor-abrogative/mimetic mutations affect mutant HTT aggregation. A Immunofluorescence of HEK293T cells overexpressing HTT Ex1 Q72-EGFP WT or phosphor-mutants (S13A/S16A, S13D/S16D, T3A, T3D, T3A/S13A/S16A, T3D/S13D/S16D) showing increased HTT aggregates in the presence of the phosphor-abrogative S13A/S16A mutation and decreased aggregation in the presence of the phosphor-mimetic S13D/S16D mutation. T3A and T3D mutations produced a comparable decrease in HTT aggregation. Expression of a triple phosphor-abrogative mutant T3A/S13A/S16A resulted in increased aggregation to a comparable degree to the double S13A/S16A mutation, while the respective phosphor-mimetic triple mutant (T3D/S13D/S16D) led to the opposite result. Nuclear signal, aggregates signal and HTT levels were assessed by DAPI staining, GFP-signal acquisition and 2B7 antibody staining respectively. (Representative images of n=3 independent experiments. Scale bars represent 50 μm). B Quantitative analysis of images in A. (Means and SDs were calculated on 3 biological replica. Oneway analysis of variance, Dunnett’s Test (*P < 0.05; **P < 0.001; ***P < 0.0001)).

**Figure 7.**
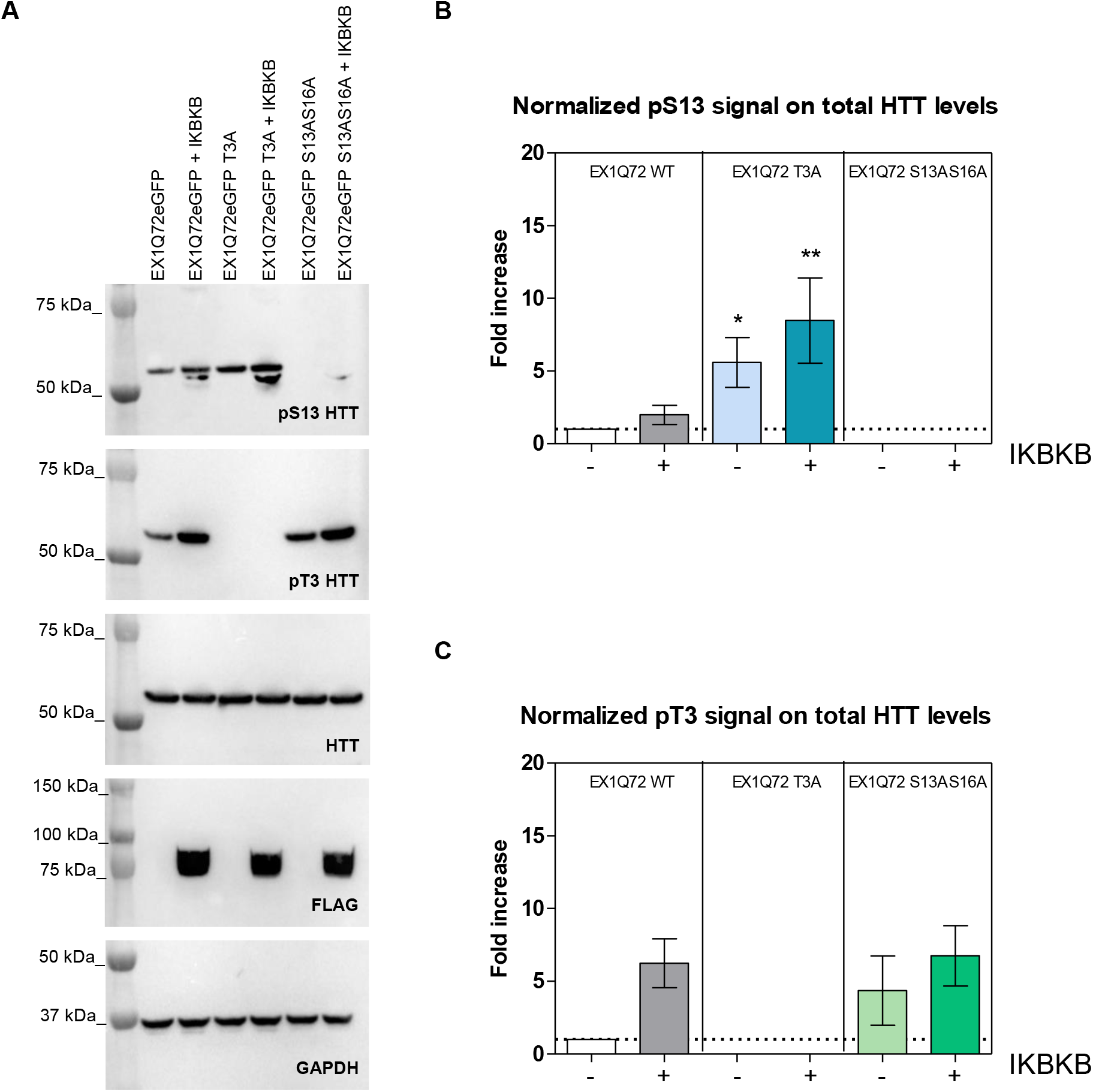
IKBKB influences pS13 HTT levels through cross-talk between T3 and S13. A Western blotting of HEK293T cells overexpressing HTT Ex1 Q72-EGFP WT or phosphor-abrogative mutants (T3A or S13A/S16A) with or without IKBKB. Protein loading, pS13 HTT levels, pT3 HTT levels, IKBKB expression and HTT levels were assessed using anti-GAPDH, anti-pS13, anti-pT3, anti-FLAG and 4C9 antibodies respectively. B-C Densitometry analysis of Western blotting in (A) showing normalized pS13 HTT levels on total HTT (B) and normalized pT3 HTT levels on total HTT (C), demonstrating that the presence of T3A mutation increases basal pS13 HTT levels and enhances the effects of IKBKB on pS13 HTT levels. (Means and SDs were calculated on 3 biological replica. One-way analysis of variance, Dunnett’s Test (*P < 0.05; **P < 0.001)).

**Figure 8.**
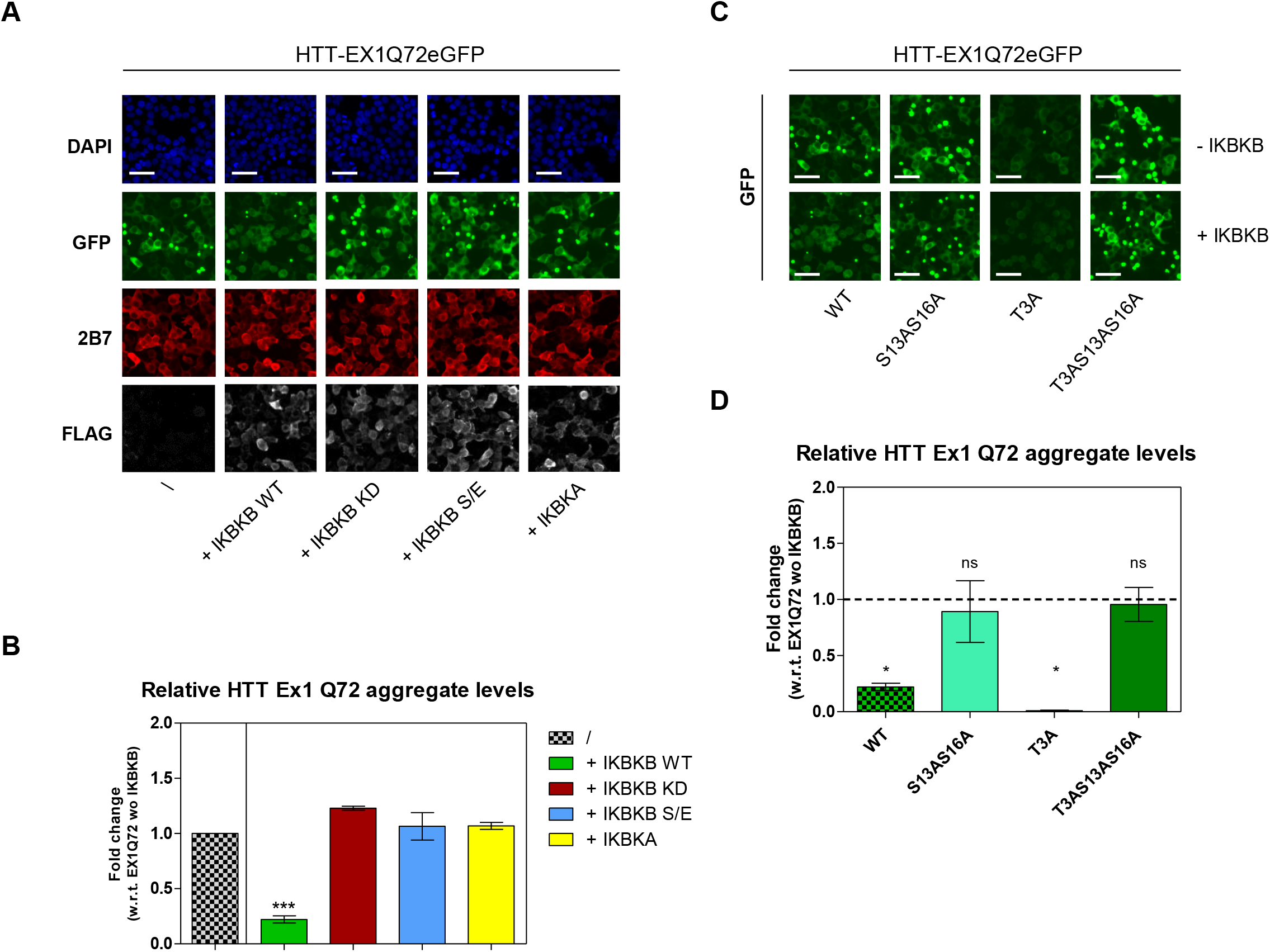
IKBKB influences mutant HTT aggregation dependent on an intact catalytic site and on the presence of S13/S16 residues. A-B HEK293T cells overexpressing HTT Ex1 Q72-EGFP WT and IKBKB WT or IKBKB KD, IKBKB S/E or IKBKA. (A) Immunofluorescence showing decreased mutant HTT aggregation only in the presence of WT IKBKB kinase. Aggregation levels are not modulated in the presence of the KD mutation, S/E mutation or in the presence of IKBKA. Nuclear signal, aggregates signal, HTT levels and kinases expression were assessed by DAPI staining, GFP-signal acquisition, 2B7 antibody and anti-FLAG antibody staining respectively. (Representative images of n=3 independent experiments. Scale bars represent 50 μm). (B) Quantitative analysis of images in (A). (Means and SDs were calculated on 3 biological replica. One-way analysis of variance, Dunnett’s Test (***P < 0.0001)). C-D HEK293T cells overexpressing HTT Ex1 Q72-EGFP WT or phosphor-abrogative mutants (S13A/S16A, T3A, T3A/S13A/S16A) with or without IKBKB. (C) Immunofluorescence showing decreased mutant HTT aggregation levels in the presence of IKBKB when S13/S16 residues are intact, with T3A modification further enhancing the capacity of IKBKB to reduce mutant HTT aggregation. Aggregate levels were assessed by GFP-signal acquisition. (Representative images of n=3 independent experiments. Scale bars represent 50 μm). (D) Quantitative analysis of images in (C). The fold change of each bar was calculated with respect to its own control without IKBKB. (Means and SDs were calculated on 3 biological replica. One-way analysis of variance, Dunnett’s Test (*P < 0.05))

## Discussion

The role of HTT PTMs in ameliorating pathological phenotypes in HD mouse models is well supported by the findings that phosphor-mimetic mutations at residues associated with phosphorylation (e.g. S13/S16 and S421) can significantly affect mutant HTT biology (Gu et al., 2009, Kratter, Zahed et al., 2016, Xu, Ng et al., 2020). Recent discoveries of pharmacological modulators of kinase activity (Ferguson & Gray, 2018, Simpson, Hughes et al., 2009), open the possibility of developing candidate tools to better increase our understanding of the mechanisms involved in modulating HTT phosphorylation and their contribution to HD pathophysiology.

However interesting conceptually, detailed studies assessing their potential therapeutic relevance has been hampered by the lack of specific and sensitive technologies to adequately determine the biophysical properties and biologically dynamic nature of bona fide PTMs. Where pharmacological modulators have been identified (Alpaugh et al., 2017, Bowie et al., 2018, Marion, Urs et al., 2014), their efficacy, potency, selectivity, precise mechanism of action and amenability as tools for further proof-of-concept studies in vivo remain to be demonstrated. As for investigational tools, several enzymes have been reported as modulators, and specifically IKBKB, TBK1 and CK2 for T3 and S13/S16 phosphorylation (Atwal et al., 2011, Bowie et al., 2018, Bustamante et al., 2015, Hegde et al., 2020, Ochaba et al., 2019, Thompson et al., 2009), PP1 for T3 phosphorylation (Branco-Santos et al., 2017), and AKT and SGK for S421 phosphorylation (Humbert, Bryson et al., 2002, Rangone, Poizat et al., 2004).

In general, however, these enzymes have not been demonstrated to potently, selectively and quantitatively influence the relevant endogenous HTT PTM, largely due to the lack of suitable readouts capable of detecting a dynamic, labile protein modification which likely affects only a portion of the total cellular protein pool. The availability and characterization of specific antibodies capable of sensitively detect HTT phosphorylation events (Aiken et al., 2009, Ansaloni, Wang et al., 2014, Atwal et al., 2011, Cariulo et al., 2017, Thompson et al., 2009) and their use in the recent development of novel ultrasensitive immunoassays (Cariulo et al., 2017, Cariulo et al., 2019) represents an appropriate platform for a more in depth validation of these current candidate modulators to provide deeper insights in the role of PTMs in HD. Of the candidate tools to increase N-terminal HTT phosphorylation, IKBKB represents one of the better validated enzymes reported to be involved in regulating pS13 or pS13/pS16 levels in cellular and animal models of HD (Atwal et al., 2011, Bustamante et al., 2015, Ochaba et al., 2019, Thompson et al., 2009), including in an endogenous HTT context in mice (Ochaba et al., 2019). Canonical (IKK complex-mediated) IKK signalling appears to be a contributor to HD toxicity mechanisms (Khoshnan & Patterson, 2011) and may control *HTT* transcription (Becanovic et al., 2015). Moreover IKBKB inhibitors increase pS13 HTT levels and decrease toxicity in cellular models of HD (Atwal et al., 2011). Although these findings suggest that inhibition of canonical IKK signalling lead to decreased toxicity and lower expression of mutant HTT, conditional IKBKB knock-out worsens the phenotype in a mouse model of HD (Ochaba et al., 2019). These paradoxical findings suggest a degree of complexity which requires further investigation (Greiner & Yang, 2011), as indeed very little is known of the downstream signalling pathways leveraged by IKBKB to increase phosphorylation of the HTT N17 domain. IKBKB is a pleiotropic kinase capable of signalling via diverse pathways (Chau, Gioia et al., 2008, Hacker & Karin, 2006, Israel, 2010, Perkins, 2007, Schrofelbauer, Polley et al., 2012) and may therefore contribute both positively and negatively to HD pathology, depending on cellular context and intracellular signalling homeostasis. Indeed, the illustration of a specific IKK pathway involved in the regulation of pS13 HTT levels adds an important aspect to the understanding of biological mechanisms involved in HD. Using a recently developed anti-pS13 antibody and ultrasensitive immunoassays (Cariulo et al., 2019), we have quantitatively examined the capacity of IKBKB to increase pS13 HTT levels expressed from the endogenous *HTT* locus in a human cell line widely employed for HTT PTM studies. We thereby identified a mechanism regulating the activity of IKBKB on the pS13 HTT substrate and propose a molecular pathway whereby IKBKB phosphorylates pS13 HTT. Importantly, we found that the ability of IKBKB to modulate pS13 HTT levels was dependent on its catalytic activity as determined by both genetic (kinase domain inactivating mutation) and pharmacological (selective IKBKB inhibitor) means. As might be expected, the effects of IKBKB on pS13 HTT levels were regulated by phosphatases, such as PP2A, likely acting at the level of IKBKB rather than by dephosphorylating HTT S13, since IKBKB-induced but not basal pS13 HTT levels were unaffected by the presence of OA, PPP2CA RNAi or PPP2CA overexpression. These observations are consistent with the described inhibitory effect of PP2A on IKBKB activity (DiDonato et al., 1997), and provide an additional level of complexity to data interpretation. Importantly, these findings offer a plausible explanation as to how pS13 HTT levels may be increased through inhibiting regulatory components of the kinase responsible for increasing pS13 HTT levels. To our knowledge, this is the first report which demonstrates a quantitative, catalytic activity-dependent, regulated modulation of endogenous HTT pS13 levels by IKBKB. The present work differentiates the IKBKB activity on pS13 HTT levels which we genetically demonstrate acts via an IKK-complex independent pathway, from IKBKB/IKK involvement in mutant HTT toxicity and *HTT* transcription, believed to act via a canonical, IKK complex-mediated pathway (Becanovic et al., 2015, Khoshnan & Patterson, 2011). The characterisation of the non-canonical, IRF-mediated IKK pathway as relevant for increasing pS13 HTT levels represents a significant advancement in the field and asks for further development of selective tools of non-canonical IKK pathway modulators to allow further research to elucidate mechanisms governing increases in pS13 HTT levels without worsening mutant HTT toxicity (Khoshnan et al., 2004) or increase its expression (Becanovic et al., 2015). We further elucidated the mechanism by which IKBKB phosphorylates HTT, providing the first evidence that IKBKB interacts directly with HTT, and that this interaction is dependent on the catalytic activity of IKBKB but independent of IKBKB’s HTT substrate phosphorylation. Finally, as a first piece of evidence that genetic modulation of pS13 HTT levels can reverse a pathologically relevant phenotype, we demonstrate, again for the first time, that increased pS13 HTT levels reduce mutant HTT aggregation in cells, fully phenocopying the effects of phosphor-mimetic mutations.

In conclusion, we believe these findings provide a significant advance in the field in several ways. From a technical perspective, this is the first demonstration that it is possible to quantitatively measure and modulate endogenous levels of a pathologically relevant PTM in HTT, phenocopying the effects of phosphor-mimetic mutations on a disease relevant readout. Conceptually, our findings advance the understanding of the relevance of IKBKB in HD biology in several ways. First, through the identification of a regulatory, phosphatase-dependent, mechanism acting via IKBKB and resulting in increased pS13 HTT levels. Second, through the identification of non-canonical, IRF-mediated IKK signalling pathway, resulting in increased pS13 HTT levels distinct from the canonical, IKK complex-mediated pathway, whose activity contributes to HD and to regulation of HTT expression. These data provide a rationale for identifying selective, non-canonical IKK signalling modulators and associated pharmacodynamic readouts, to further elucidate candidate pharmacological tools and pathways involved in HD pathophysiology.

## Materials and Methods

### Antibodies

The MW1, 2B7 and 4C9 antibodies bind to the polyQ stretch, the N17 domain and the polyproline region of HTT respectively. They were obtained from the CHDI Foundation (New York, NY) and their use in Western blotting and SMC assays has been described previously (Cariulo et al., 2017). The anti-pS13 antibody is an affinity purified, rabbit polyclonal antibody specific for the phosphorylated S13 residue of human huntingtin raised against the peptide LMKAFE(pS)LKSFQ developed by the CHDI foundation and available from the Coriell Institute for Biomedical research, HD Community Biorepository (ID CH01115). The validation of this antibody in different applications has been formerly reported (Cariulo et al., 2019). The here used anti-pT3 antibody was previously described (Cariulo et al., 2017). Monoclonal antibodies mAb2166 (catalog #MAB2166; Merck) and anti-GAPDH (catalog #G9545; Sigma-Aldrich), anti-FLAG mouse antibody (catalog #F1804; Sigma-Aldrich), anti-FLAG rabbit antibody (catalog #F7425; Sigma Aldrich), anti-IKBKB antibody (catalog #ab32135; Abcam), anti-pS176/180-IKBKB antibody (catalog #2697; Cell Signaling), Anti-Glial Fibrillary Acidic Protein GFAP (catalog #G9269; Sigma-Aldrich), anti-PP2A antibody (catalog #SAB4200266; Sigma-Aldrich), anti-IRF3 antibody (catalog #AB76409; Abcam) and anti-pS386-IRF3 antibody (catalog #AB76493; Abcam) were all supplied by a commercial source. Secondary antibodies used for Western blotting were Goat-Anti-mouse IgG HRP conjugated (catalog #12-349; Merck) and Goat-Anti-rabbit IgG HRP conjugated (catalog #12-348; Merck). The D2-fluophore and terbium antibody labelling for TR-FRET assay were custom made at CisBio (Bagnols, France). The Alexa-647 labelling for detection used in SMC and in ELISA assays was performed using the Alexa Fluor-647 Monoclonal Antibody Labeling Kit from Thermo Fisher Scientific (catalog #A20186) following manufacturer’s instructions. MW1 antibody was conjugated to magnetic particles for SMC assays, following the manufacturer’s recommendations (catalog #03-0077-02; Merck).

### Plasmid and constructs

All plasmids employed in this study have been ordered or custom synthetized at TEMA Ricerca or Genescript (Nanjing, China) respectively. The chosen vector for constructs IKBKB, IKBKB KD, IKBKB S/E, IKBKB NBD, IKBKB LZ is pCDNA3.1. IKBKA (#RC216718), IKBKG (#RC218044), IKBKE (#RC212481) and PPP2CA (#RC201334) are commercially sourced by TEMA Ricerca. These constructs are also FLAG-tagged at the C-terminus. cDNAs constructs encoding for N-terminal HTT fragments (exon 1 HTT 1-90, based on Q23 numbering; Table S2) bearing different polyQ lengths (Q16 or Q72) and mutations (T3A, S13A/S16A, T3D, S13D/S16D, T3A/S13A/S16A, T3D/S13D/S16D) codify also for a EGFP-Tag at the C-terminus and are inserted in a pCMV vector. N-terminal HTT fragment, HTT 1-571, based on Q23 numbering (Table S3), is encoded in an untagged pCDNA3.1 vector. Expression of all constructs in mammalian cells has been validated as previously reported (Cariulo et al., 2019, Fodale et al., 2014).

### HEK293T cell culture and manipulation

For Western blotting and immunoassay analysis HEK293T cells were cultured, transfected and lysed as previously described (Cariulo et al., 2019). For immunofluorescence analysis, HEK293T cells were plated in Poly-L-lysine (100 μg/ml) coated 96-well black plates with μClear bottom (catalog #655090; Greiner) at 20.000 cells per well and transfected using Lipofectamine 2000 (Thermo Fisher Scientific) according to manufacturer’s protocols. Lysis was performed 24 hours from transfection in lysis buffer (TBS, 0.4% Triton X100) supplemented with 1X protease inhibitor cocktail (catalog #11697498001; Roche) and phosphatase inhibitor cocktail (catalog #04986837001; Roche). Okadaic Acid (catalog #5934S; Cell Signaling) was delivered to HEK293T cells at a concentration of 350 nM for 1 hour at 37°C before their lysis. The IKBKB inhibitor compound (Bay-65-1942, (Atwal et al., 2011)) was sourced by the CHDI foundation and HEK293T cells were treated for 24 hours at 37°C before lysis. The combination of the treatment with OA and Bay-65-1942 was performed as follows: HEK293T cells were transfected as previously reported; after 24 hours from transfection, Bay-65-1942 compound was delivered to cells at the reported concentration, medium was changed and treatment was carried out for 24 hours; 1 hour before lysis, cells were treated with OA as previously described. Silencing via RNA interference (RNAi) was performed using specific PP2A-Cα siRNA commercially supplied (catalog #sc-43509; Santa Cruz Biotechnology). The siRNA transfection was performed as reported above, employing the siRNAs at the final concentration of 100 nM. Lysis of cells was carried out after 48 hours from transfection.

### Western blot

Samples were denatured at 95°C in 4X Loading Buffer (125 mM Tris-HCl pH 6.8, 6% SDS, 4 M urea, 4 mM EDTA, 30% Glycerol, 4% 2-Mercaptoethanol and Bromophenol Blue) and loaded on NuPAGE 4-12% Bis-Tris Gel (catalog #WG1402BOX; Thermo Fisher Scientific). Proteins were transferred on PVDF membrane (catalog #162-0177; Bio-Rad Laboratories) using wet blotting. After fixing in 0.4% paraformaldehyde solution and blocking with 5% non-fat milk in TBS/0.1% Tween-20, primary antibodies incubation was carried out overnight at 4°C and secondary antibody incubations for 1 hour at room temperature. Protein bands were detected using chemiluminescent substrate (Supersignal West Femto Maximum catalog #3406; Supersignal West Pico Maximum catalog #34087; Thermo Fisher Scientific) on Chemidoc XRS+ (Bio-Rad Laboratories). Densitometric analysis was performed using ImageJ software.

### Immunoprecipitation

Immunoprecipitation was performed using Dynabeads^®^ Protein G (catalog #10004D; Thermo Fisher Scientific) following the manufacturer’s instructions and using anti-FLAG (mouse) antibody or an unrelated antibody (Anti-Glial Fibrillary Acidic Protein GFAP) as previously described (Vezzoli, Caron et al., 2019). The pulled down material was loaded on a SDS-PAGE and the Western Blot was performed as described above.

### SMC assay

50 μL/well of dilution buffer (6% BSA, 0.8% Triton X-100, 750 mM NaCl, and protease inhibitor cocktail) were added to a 96-well plate (catalog #P-96-450V-C; Axygen). Samples to be tested were diluted in artificial cerebral spinal fluid (ACSF: 0.3 M NaCl; 6 mM KCl; 2.8 mM CaCl2-2H2O; 1.6 mM MgCl2-6H20; 1.6 mM Na2HPO4-7H2O; 0.4 mM NaH2PO4-H2O) supplemented with 1% Tween-20 and protease inhibitor cocktail, in a final volume of 150 μL/well. Finally, 100 μL/well of the MW1 antibody coupled with magnetic particles (appropriately diluted in Erenna Assay buffer catalog #02-0474-00; Merck), were added to the assay plate, and incubated for 1 hour at room temperature under orbital shaking for the capturing step. The beads were then washed with Erenna System buffer (catalog #02-0111-00; Merck) and resuspended using 20 μL/well of the specific detection antibody labeled with Alexa-647 fluorophore appropriately diluted in Erenna Assay buffer (anti-pS13 antibody for pS13 HTT levels readout and 2B7 for total HTT levels readout). The plate was incubated for 1 hour at room temperature under shaking. After washing, the beads were resuspended and transferred in a new 96-well plate. 10 μL/well of Erenna buffer B (catalog #02-0297-00; Merck) were added to the beads for elution and incubated for 5 minutes at room temperature under orbital shaking. The eluted complex was magnetically separated from the beads and transferred in a 384-well plate (Nunc catalog #264573; Sigma) where it was neutralized with 10 μL/well of Erenna buffer D (catalog #02-0368-00; Merck). Finally, the 384-well plate was heat-sealed and analyzed with the Erenna Immunoassay System.

### TR-FRET assay

5 μL/well of samples and 1 μL/well of antibodies cocktail (2B7-Tb 1 ng/μL; FLAG-D2 10 ng/μL) diluted in lysis buffer (composition described before) were added to the 384-well assay plate (Low Volume-F Bottom catalog #784080; Greiner). Assay was performed according to (Fodale et al., 2014).

### ELISA assays

Assay plates (Nunc^™^ MaxiSorp^™^ 96-well polystyrene plate catalog #449824; Thermo Fisher Scientific) were coated with 100 μL/well of the capture antibody (2B7 or anti-PP2A) diluted in TBS at the final concentration of 1 μg/mL and incubated without shaking overnight at 4°C. After washing the plate three times with wash buffer (TBS, 0.1% Tween20), the plate was blocked with 300 μL/well of BSA 1% in TBS (Blocking Buffer) for 30 minutes at room temperature and washed three times with wash buffer. 100 μL/well of the analyte diluted at the appropriate concentration in sample buffer (ACSF supplemented with 1% Tween-20 and protease inhibitor cocktail) was added to the wells and incubated for 1 hour at room temperature. After three washing steps, 50 μL/well of detection antibody (anti-FLAG-D2) diluted in Erenna Assay buffer (catalog #02-0474-00; Merck) at the final concentration of 1 μL/mL were added to the assay plate and incubated for 1 hour at room temperature. After five washing steps, 20 μL/well of Erenna buffer B (catalog #02-0297-00; Merck) were added to the assay plate and incubated for 5 minutes at room temperature under orbital shaking. 20 μL/well of the neutralization Erenna buffer D (catalog #02-0368-00; Merck) were added to the assay plate. After 5 minutes of incubation at room temperature under mild agitation, 35 μL/well of the neutralized samples were transferred in a final 384-well plate (Nunc catalog #264573; Sigma). The 384-well plate was heat-sealed and analyzed with the Erenna Immunoassay System.

### High-content imaging and analysis

After 24 hours from transfection (described before), medium was removed and HEK293T cells were washed 3 times with TBS and fixed for 20 min at room-temperature in 4% paraformaldehyde/4% sucrose in TBS solution. After two washes in TBS, cells were incubated with blocking solution (1% Donkey Serum, 0.1% Triton-X, 3% BSA in TBS) for 1 hour at room temperature and then incubated with primary 2B7 and anti-FLAG (rabbit) antibodies (previously described) diluted 1:300 in staining solution (0.2% Triton-X, 3% BSA in TBS) overnight at 4 °C under constant mild agitation. The next day cells were washed 3 times in TBS and incubated for 1 hour at room temperature with donkey anti-mouse IgG antibody conjugated to Alexa-647 or donkey anti-rabbit IgG antibody conjugated to Alexa-555 (1:2000 in staining solution) provided by a commercial source (catalog #A31571 and catalog #A31572; Life Technologies) together with 1 μg/ml of Hoechst staining solution (catalog #H33342; Sigma-Aldrich). After 3 washes with TBS, images were acquired in an automated fashion employing the Incell-2000 high-content imaging system (GE-Healthcare) using a 20x objective and appropriate filters. Three independent experiments were performed. For each experiment every condition was evaluated in technical duplicates, 2 well/conditions and 5 fields per well were acquired. Imaging times settings were: 50 ms for H33342 (DAPI), 30 ms for GFP and 700 ms for Alexa-647 and Alexa-555 (2B7 and anti-FLAG). Images were analyzed using the Developer toolbox 1.9.1 software (GE-Healthcare). For counting of aggregates, a vesicle algorithm was employed identifying aggregates based on shape, size and intensity of small objects.

### SMC data analysis and statistics

Normalized pS13 signal on total HTT levels measured by SMC assay was determined as the ratio between the fold increase obtained from MW1/pS13 SMC signal (pS13 HTT levels readout) and the fold increase calculated on the MW1/2B7 SMC signal (total HTT levels readout). This ratio was derived as follows: MW1/pS13SMC assay and MW1/2B7 SMC assay were performed in parallel on the same samples which were analyzed in a serial dilution curve (6 dilution points 1:3 plus blank, technical duplicates). For each readout, a curve fitting per sample (described by a four-parameter logistic curve fit) was calculated, and a fold increase among curves (which shared the top, the bottom and the slope) was assessed basing on EC50 parameter (fixing as a reference one of the analyzed samples). The statistical significance was assessed using a paired t-test (two-tailed) in the case of the analysis of n=2 samples, or using a One-way analysis of variance, applying Dunnett’s Test (which allows to compare all columns versus a fix control column) where n > 2 samples need to be compared. Graphs were generated and statistical analysis was performed using the software GraphPad Prism 6.

## Acknowledgements

The work was funded by CHDI Foundation Inc., a non-profit biomedical research organization exclusively dedicated to collaboratively developing therapeutics that will substantially improve the lives of individuals with Huntington’s disease.

## Author contributions

CC, PM and MV: Investigation, formal analysis and data curation

RL, LTS and CD: Funding acquisition, resources and supervision

CC, LP and AC: Conceptualization and writing - original draft

LP and AC: Project administration

## Figure legends

**Figure S1.**
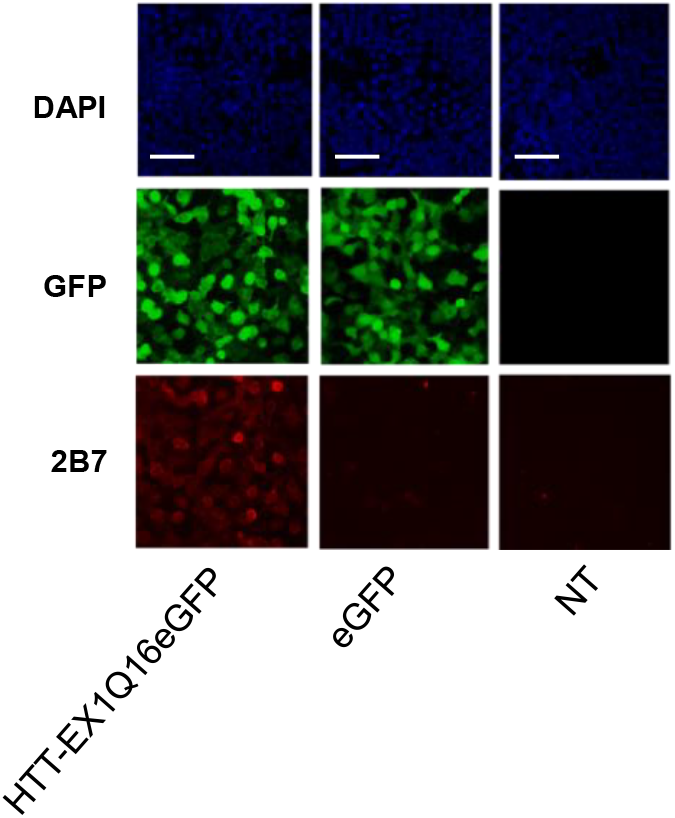
Negative controls for immunofluorescence analysis. Immunofluorescence of HEK293T cells untransfected (NT) or transfected with eGFP or Ex1 Q16-EGFP constructs reported as negative controls for high-content imaging analysis. The overexpression of Ex1 Q16-EGFP is properly detected by 2B7 antibody staining as well as GFP-signal acquisition, without showing aggregates signal. Nuclear signal was assessed by DAPI staining. (Representative images of n=3 independent experiments. Scale bars represent 50 μm).

**Table S2.**
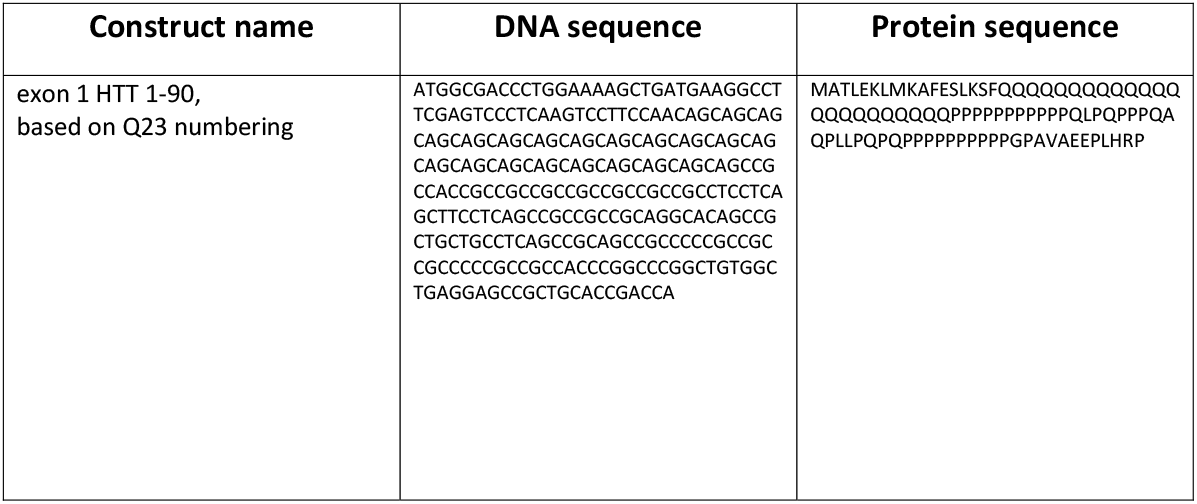
DNA and amino acid sequence of HTT-exon1 construct

**Table S3.**
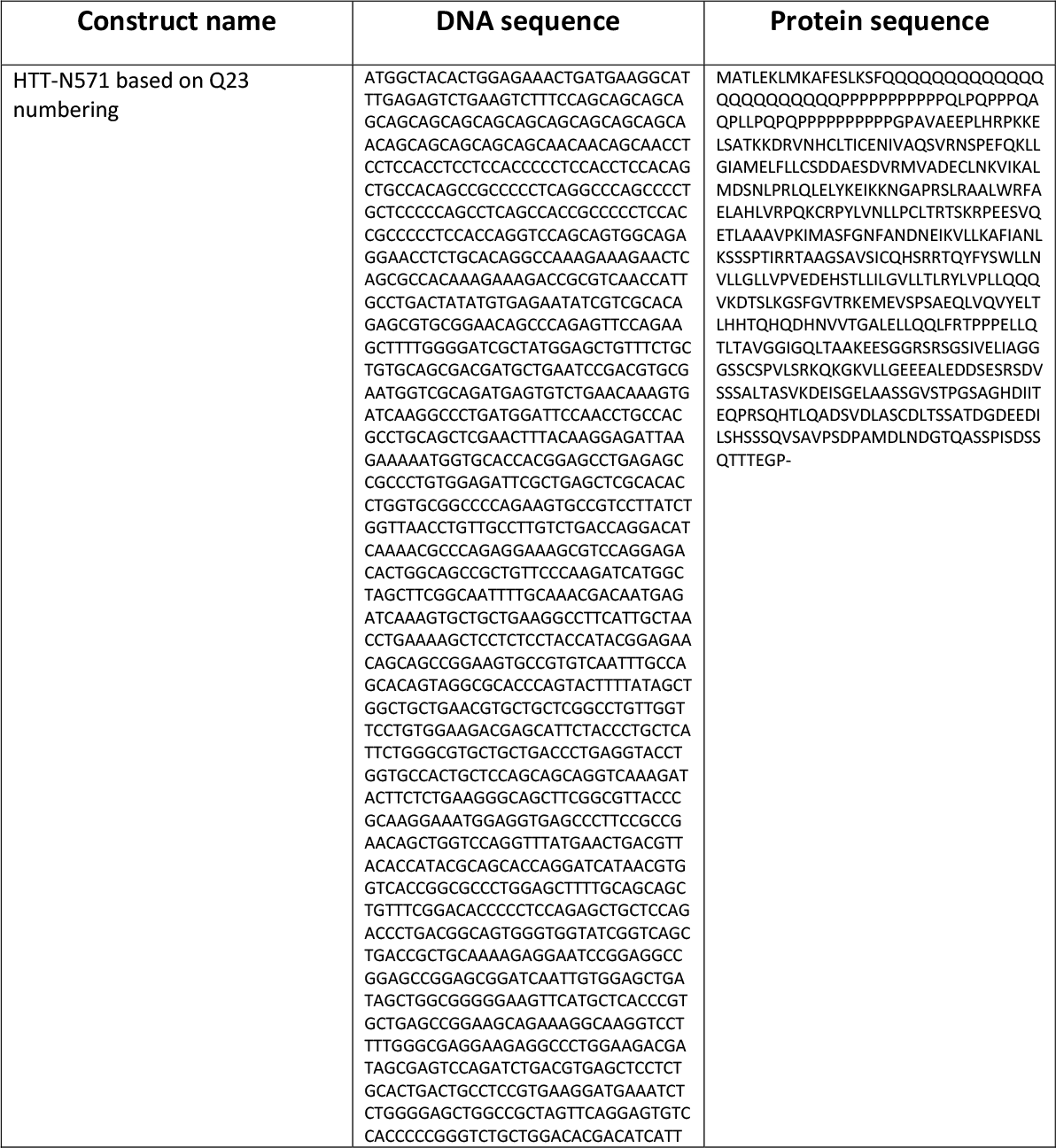
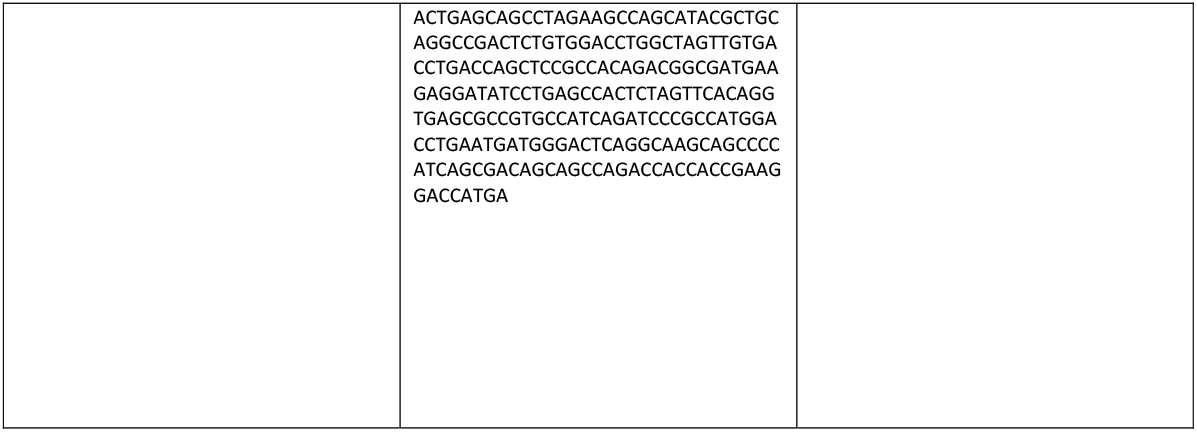
DNA and amino acid sequence of HTT-N571 construct

## References

Aiken CT, Steffan JS, Guerrero CM, Khashwji H, Lukacsovich T, Simmons D, Purcell JM, Menhaji K, Zhu YZ, Green K, Laferla F, Huang L, Thompson LM, Marsh JL (2009) Phosphorylation of threonine 3: implications for Huntingtin aggregation and neurotoxicity. J Biol Chem 284: 29427–36

Alpaugh M, Galleguillos D, Forero J, Morales LC, Lackey SW, Kar P, Di Pardo A, Holt A, Kerr BJ, Todd KG, Baker GB, Fouad K, Sipione S (2017) Disease-modifying effects of ganglioside GM1 in Huntington’s disease models. EMBO Mol Med 9: 1537–1557

Ansaloni A, Wang ZM, Jeong JS, Ruggeri FS, Dietler G, Lashuel HA (2014) One-pot semisynthesis of exon 1 of the Huntingtin protein: new tools for elucidating the role of posttranslational modifications in the pathogenesis of Huntington’s disease. Angew Chem Int Ed Engl 53: 1928–33

Atwal RS, Desmond CR, Caron N, Maiuri T, Xia J, Sipione S, Truant R (2011) Kinase inhibitors modulate huntingtin cell localization and toxicity. Nat Chem Biol 7: 453–60

Atwal RS, Xia J, Pinchev D, Taylor J, Epand RM, Truant R (2007) Huntingtin has a membrane association signal that can modulate >huntingtin aggregation, nuclear entry and toxicity. Hum Mol Genet 16: 2600–15

Balka KR, Louis C, Saunders TL, Smith AM, Calleja DJ, D’Silva DB, Moghaddas F, Tailler M, Lawlor KE, Zhan Y, Burns CJ, Wicks IP, Miner JJ, Kile BT, Masters SL, De Nardo D (2020) TBK1 and IKKepsilon Act Redundantly to Mediate STING-Induced NF-kappaB Responses in Myeloid Cells. Cell Rep 31: 107492

Barisic S, Strozyk E, Peters N, Walczak H, Kulms D (2008) Identification of PP2A as a crucial regulator of the NF-kappaB feedback loop: its inhibition by UVB turns NF-kappaB into a pro-apoptotic factor. Cell Death Differ 15: 1681–90

Bates G (2003) Huntingtin aggregation and toxicity in Huntington’s disease. Lancet 361: 1642–4

Bates GP, Dorsey R, Gusella JF, Hayden MR, Kay C, Leavitt BR, Nance M, Ross CA, Scahill RI, Wetzel R, Wild EJ, Tabrizi SJ (2015) Huntington disease. Nature Reviews Disease Primers: 15005

Becanovic K, Norremolle A, Neal SJ, Kay C, Collins JA, Arenillas D, Lilja T, Gaudenzi G, Manoharan S, Doty CN, Beck J, Lahiri N, Portales-Casamar E, Warby SC, Connolly C, De Souza RA, Network RIotEHsD, Tabrizi SJ, Hermanson O, Langbehn DR et al. (2015) A SNP in the HTT promoter alters NF-kappaB binding and is a bidirectional genetic modifier of Huntington disease. Nat Neurosci 18: 807–16

Bowie LE, Maiuri T, Alpaugh M, Gabriel M, Arbez N, Galleguillos D, Hung CLK, Patel S, Xia J, Hertz NT, Ross CA, Litchfield DW, Sipione S, Truant R (2018) N6-Furfuryladenine is protective in Huntington’s disease models by signaling huntingtin phosphorylation. Proc Natl Acad Sci U S A 115: E7081–E7090

Branco-Santos J, Herrera F, Pocas GM, Pires-Afonso Y, Giorgini F, Domingos PM, Outeiro TF (2017) Protein phosphatase 1 regulates huntingtin exon 1 aggregation and toxicity. Hum Mol Genet 26: 3763–3775

Bustamante MB, Ansaloni A, Pedersen JF, Azzollini L, Cariulo C, Wang ZM, Petricca L, Verani M, Puglisi F, Park H, Lashuel H, Caricasole A (2015) Detection of huntingtin exon 1 phosphorylation by Phos-Tag SDS-PAGE: Predominant phosphorylation on threonine 3 and regulation by IKKbeta. Biochem Biophys Res Commun 463: 1317–22

Cariulo C, Azzollini L, Verani M, Martufi P, Boggio R, Chiki A, Deguire SM, Cherubini M, Gines S, Marsh JL, Conforti P, Cattaneo E, Santimone I, Squitieri F, Lashuel HA, Petricca L, Caricasole A (2017) Phosphorylation of huntingtin at residue T3 is decreased in Huntington’s disease and modulates mutant huntingtin protein conformation. Proc Natl Acad Sci U S A 114: E10809–E10818

Cariulo C, Verani M, Martufi P, Ingenito R, Finotto M, Deguire SM, Lavery DJ, Toledo-Sherman L, Lee R, Doherty EM, Vogt TF, Dominguez C, Lashuel HA, Petricca L, Caricasole A (2019) Ultrasensitive quantitative measurement of huntingtin phosphorylation at residue S13. Biochem Biophys Res Commun

Chatterjee M, Steffan JS, Lukacsovich T, Marsh JL, Agrawal N (2021) Serine residues 13 and 16 are key modulators of mutant huntingtin induced toxicity in Drosophila. Exp Neurol 338: 113463

Chau TL, Gioia R, Gatot JS, Patrascu F, Carpentier I, Chapelle JP, O’Neill L, Beyaert R, Piette J, Chariot A (2008) Are the IKKs and IKK-related kinases TBK1 and IKK-epsilon similarly activated? Trends Biochem Sci 33: 171–80

Chiki A, Ricci J, Hegde R, Abriata LA, Reif A, Boudeffa D, Lashuel HA (2021) Site-Specific Phosphorylation of Huntingtin Exon 1 Recombinant Proteins Enabled by the Discovery of Novel Kinases. Chembiochem 22: 217–231

Chiki A. DSM, Ruggeri F.S., Cendrowska U., Ansaloni A., Wang Z.M., Sanfelice D., Burai R., Vieweg S., Pastore P., Dietler G., Lashuel H.A. (2017) Huntingtin structure and aggregation is regulated by cross-talk between N-terminal post-translational modifications. Angew Chem Int Ed Engl Submitted

Clark K, Peggie M, Plater L, Sorcek RJ, Young ER, Madwed JB, Hough J, McIver EG, Cohen P (2011) Novel cross-talk within the IKK family controls innate immunity. Biochem J 434: 93–104

Cong X, Held JM, DeGiacomo F, Bonner A, Chen JM, Schilling B, Czerwieniec GA, Gibson BW, Ellerby LM (2011) Mass spectrometric identification of novel lysine acetylation sites in huntingtin. Mol Cell Proteomics 10: M111 009829

Cooper JK, Schilling G, Peters MF, Herring WJ, Sharp AH, Kaminsky Z, Masone J, Khan FA, Delanoy M, Borchelt DR, Dawson VL, Dawson TM, Ross CA (1998) Truncated N-terminal fragments of huntingtin with expanded glutamine repeats form nuclear and cytoplasmic aggregates in cell culture. Hum Mol Genet 7: 783–90

Crick SL, Ruff KM, Garai K, Frieden C, Pappu RV (2013) Unmasking the roles of N- and C-terminal flanking sequences from exon 1 of huntingtin as modulators of polyglutamine aggregation. Proc Natl Acad Sci U S A 110: 20075–80

Daldin M, Fodale V, Cariulo C, Azzollini L, Verani M, Martufi P, Spiezia MC, Deguire SM, Cherubini M, Macdonald D, Weiss A, Bresciani A, Vonsattel JG, Petricca L, Marsh JL, Gines S, Santimone I, Marano M, Lashuel HA, Squitieri F et al. (2017) Polyglutamine expansion affects huntingtin conformation in multiple Huntington’s disease models. Sci Rep 7: 5070

DeGuire S.M. RFS, Fares M.B., Cendrowska U., Dietler G., Lashuel H.A. (2017) Dissecting the role of N-terminal serine phosphorylation. Hum Mol Genet Submitted

DeGuire SM, Ruggeri FS, Fares MB, Chiki A, Cendrowska U, Dietler G, Lashuel HA (2018) N-terminal Huntingtin (Htt) phosphorylation is a molecular switch regulating Htt aggregation, helical conformation, internalization, and nuclear targeting. J Biol Chem 293: 18540–18558

Di Pardo A, Maglione V, Alpaugh M, Horkey M, Atwal RS, Sassone J, Ciammola A, Steffan JS, Fouad K, Truant R, Sipione S (2012) Ganglioside GM1 induces phosphorylation of mutant huntingtin and restores normal motor behavior in Huntington disease mice. Proc Natl Acad Sci U S A 109: 3528–33

DiDonato JA, Hayakawa M, Rothwarf DM, Zandi E, Karin M (1997) A cytokine-responsive IkappaB kinase that activates the transcription factor NF-kappaB. Nature 388: 548–54

Dounay AB, Forsyth CJ (2002) Okadaic acid: the archetypal serine/threonine protein phosphatase inhibitor. Curr Med Chem 9: 1939–80

Ehrnhoefer DE, Sutton L, Hayden MR (2011) Small changes, big impact: posttranslational modifications and function of huntingtin in Huntington disease. Neuroscientist 17: 475–92

Ferguson FM, Gray NS (2018) Kinase inhibitors: the road ahead. Nat Rev Drug Discov 17: 353–377

Fitzgerald KA, McWhirter SM, Faia KL, Rowe DC, Latz E, Golenbock DT, Coyle AJ, Liao SM, Maniatis T (2003) IKKepsilon and TBK1 are essential components of the IRF3 signaling pathway. Nat Immunol 4: 491–6

Flower M, Lomeikaite V, Ciosi M, Cumming S, Morales F, Lo K, Hensman Moss D, Jones L, Holmans P, Investigators T-H, Consortium O, Monckton DG, Tabrizi SJ (2019) MSH3 modifies somatic instability and disease severity in Huntington’s and myotonic dystrophy type 1. Brain>

Fodale V, Boggio R, Daldin M, Cariulo C, Spiezia MC, Byrne LM, Leavitt BR, Wild EJ, Macdonald D, Weiss A, Bresciani A (2017) Validation of Ultrasensitive Mutant Huntingtin Detection in Human Cerebrospinal Fluid by Single Molecule Counting Immunoassay. J Huntingtons Dis 6: 349–361

Fodale V, Kegulian NC, Verani M, Cariulo C, Azzollini L, Petricca L, Daldin M, Boggio R, Padova A, Kuhn R, Pacifici R, Macdonald D, Schoenfeld RC, Park H, Isas JM, Langen R, Weiss A, Caricasole A (2014) Polyglutamine-and temperature-dependent conformational rigidity in mutant huntingtin revealed by immunoassays and circular dichroism spectroscopy. PLoS One 9: e112262

Franich NR, Hickey MA, Zhu C, Osborne GF, Ali N, Chu T, Bove NH, Lemesre V, Lerner RP, Zeitlin SO, Howland D, Neueder A, Landles C, Bates GP, Chesselet MF (2019) Phenotype onset in Huntington’s disease knock-in mice is correlated with the incomplete splicing of the mutant huntingtin gene. J Neurosci Res 97: 1590–1605

Genetic Modifiers of Huntington’s Disease Consortium. Electronic address ghmhe, Genetic Modifiers of Huntington’s Disease C (2019) CAG Repeat Not Polyglutamine Length Determines Timing of Huntington’s Disease Onset. Cell 178: 887–900 e14

Gipson TA, Neueder A, Wexler NS, Bates GP, Housman D (2013) Aberrantly spliced HTT, a new player in Huntington’s disease pathogenesis. RNA Biol 10: 1647–52

Goold R, Flower M, Moss DH, Medway C, Wood-Kaczmar A, Andre R, Farshim P, Bates GP, Holmans P, Jones L, Tabrizi SJ (2019) FAN1 modifies Huntington’s disease progression by stabilizing the expanded HTT CAG repeat. Hum Mol Genet 28: 650–661

Greiner ER, Yang XW (2011) Huntington’s disease: flipping a switch on huntingtin. Nat Chem Biol 7: 412–4

Gu X, Cantle JP, Greiner ER, Lee CY, Barth AM, Gao F, Park CS, Zhang Z, Sandoval-Miller S, Zhang RL, Diamond M, Mody I, Coppola G, Yang XW (2015) N17 Modifies mutant Huntingtin nuclear pathogenesis and severity of disease in HD BAC transgenic mice. Neuron 85: 726–41

Gu X, Greiner ER, Mishra R, Kodali R, Osmand A, Finkbeiner S, Steffan JS, Thompson LM, Wetzel R, Yang XW (2009) Serines 13 and 16 are critical determinants of full-length human mutant huntingtin induced disease pathogenesis in HD mice. Neuron 64: 828–40

Hacker H, Karin M (2006) Regulation and function of IKK and IKK-related kinases. Sci STKE 2006: re13

Hauenstein AV, Rogers WE, Shaul JD, Huang DB, Ghosh G, Huxford T (2014) Probing kinase activation and substrate specificity with an engineered monomeric IKK2. Biochemistry 53: 2064–73

Havel LS, Wang CE, Wade B, Huang B, Li S, Li XJ (2011) Preferential accumulation of N-terminal mutant huntingtin in the nuclei of striatal neurons is regulated by phosphorylation. Hum Mol Genet 20: 1424–37

Hegde RN, Chiki A, Petricca L, Martufi P, Arbez N, Mouchiroud L, Auwerx J, Landles C, Bates GP, Singh-Bains MK, Dragunow M, Curtis MA, Faull RL, Ross CA, Caricasole A, Lashuel HA (2020) TBK1 phosphorylates mutant Huntingtin and suppresses its aggregation and toxicity in Huntington’s disease models. EMBO J 39: e104671

Holmans PA, Massey TH, Jones L (2017) Genetic modifiers of Mendelian disease: Huntington’s disease and the trinucleotide repeat disorders. Hum Mol Genet 26: R83–R90

Huang B, Lucas T, Kueppers C, Dong X, Krause M, Bepperling A, Buchner J, Voshol H, Weiss A, Gerrits B, Kochanek S (2015) Scalable production in human cells and biochemical characterization of full-length normal and mutant huntingtin. PLoS One 10: e0121055

Humbert S, Bryson EA, Cordelieres FP, Connors NC, Datta SR, Finkbeiner S, Greenberg ME, Saudou F (2002) The IGF-1/Akt pathway is neuroprotective in Huntington’s disease and involves Huntingtin phosphorylation by Akt. Dev Cell 2: 831–7

Huynh QK, Boddupalli H, Rouw SA, Koboldt CM, Hall T, Sommers C, Hauser SD, Pierce JL, Combs RG, Reitz BA, Diaz-Collier JA, Weinberg RA, Hood BL, Kilpatrick BF, Tripp CS (2000) Characterization of the recombinant IKK1/IKK2 heterodimer. Mechanisms regulating kinase activity. J Biol Chem 275: 25883–91

Israel A (2010) The IKK complex, a central regulator of NF-kappaB activation. Cold Spring Harb Perspect Biol 2: a000158

Kelley NW, Huang X, Tam S, Spiess C, Frydman J, Pande VS (2009) The predicted structure of the headpiece of the Huntingtin protein and its implications on Huntingtin aggregation. J Mol Biol 388: 919–27

Khoshnan A, Ko J, Tescu S, Brundin P, Patterson PH (2009) IKKalpha and IKKbeta regulation of DNA damage-induced cleavage of huntingtin. PLoS One 4: e5768

Khoshnan A, Ko J, Watkin EE, Paige LA, Reinhart PH, Patterson PH (2004) Activation of the IkappaB kinase complex and nuclear factor-kappaB contributes to mutant huntingtin neurotoxicity. J Neurosci 24: 7999–8008

Khoshnan A, Patterson PH (2011) The role of IkappaB kinase complex in the neurobiology of Huntington’s disease. Neurobiol Dis 43: 305–11

Khoshnan A, Sabbaugh A, Calamini B, Marinero SA, Dunn DE, Yoo JH, Ko J, Lo DC, Patterson PH (2017) IKKbeta and mutant huntingtin interactions regulate the expression of IL-34: implications for microglial-mediated neurodegeneration in HD. Hum Mol Genet 26: 4267–4277

Kishore N, Huynh QK, Mathialagan S, Hall T, Rouw S, Creely D, Lange G, Caroll J, Reitz B, Donnelly A, Boddupalli H, Combs RG, Kretzmer K, Tripp CS (2002) IKK-i and TBK-1 are enzymatically distinct from the homologous enzyme IKK-2: comparative analysis of recombinant human IKK-i, TBK-1, and IKK-2. J Biol Chem 277: 13840–7

Kratter IH, Zahed H, Lau A, Tsvetkov AS, Daub AC, Weiberth KF, Gu X, Saudou F, Humbert S, Yang XW, Osmand A, Steffan JS, Masliah E, Finkbeiner S (2016) Serine 421 regulates mutant huntingtin toxicity and clearance in mice. J Clin Invest 126: 3585–97

Landles C, Sathasivam K, Weiss A, Woodman B, Moffitt H, Finkbeiner S, Sun B, Gafni J, Ellerby LM, Trottier Y, Richards WG, Osmand A, Paganetti P, Bates GP (2010) Proteolysis of mutant huntingtin produces an exon 1 fragment that accumulates as an aggregated protein in neuronal nuclei in Huntington disease. J Biol Chem 285: 8808–23

Lee JM, Chao MJ, Harold D, Abu Elneel K, Gillis T, Holmans P, Jones L, Orth M, Myers RH, Kwak S, Wheeler VC, MacDonald ME, Gusella JF (2017) A modifier of Huntington’s disease onset at the MLH1 locus. Hum Mol Genet 26: 3859–3867

Li L, Liu H, Dong P, Li D, Legant WR, Grimm JB, Lavis LD, Betzig E, Tjian R, Liu Z (2016) Real-time imaging of Huntingtin aggregates diverting target search and gene transcription. Elife 5

Li SH, Li XJ (1998) Aggregation of N-terminal huntingtin is dependent on the length of its glutamine repeats. Hum Mol Genet 7: 777–82

Liu S, Misquitta YR, Olland A, Johnson MA, Kelleher KS, Kriz R, Lin LL, Stahl M, Mosyak L (2013) Crystal structure of a human IkappaB kinase beta asymmetric dimer. J Biol Chem 288: 22758–67

Maiuri T, Woloshansky T, Xia J, Truant R (2013) The huntingtin N17 domain is a multifunctional CRM1 and Ran-dependent nuclear and cilial export signal. Hum Mol Genet 22: 1383–94

Mangiarini L, Sathasivam K, Seller M, Cozens B, Harper A, Hetherington C, Lawton M, Trottier Y, Lehrach H, Davies SW, Bates GP (1996) Exon 1 of the HD gene with an expanded CAG repeat is sufficient to cause a progressive neurological phenotype in transgenic mice. Cell 87: 493–506

Marion S, Urs NM, Peterson SM, Sotnikova TD, Beaulieu JM, Gainetdinov RR, Caron MG (2014) Dopamine D2 receptor relies upon PPM/PP2C protein phosphatases to dephosphorylate huntingtin protein. J Biol Chem 289: 11715–24

May MJ, Marienfeld RB, Ghosh S (2002) Characterization of the Ikappa B-kinase NEMO binding domain. J Biol Chem 277: 45992–6000

McColgan P, Tabrizi SJ (2018) Huntington’s disease: a clinical review. Eur J Neurol 25: 24–34

Menalled L, Brunner D (2014) Animal models of Huntington’s disease for translation to the clinic: best practices. Mov Disord 29: 1375–90

Mercurio F, Zhu H, Murray BW, Shevchenko A, Bennett BL, Li J, Young DB, Barbosa M, Mann M, Manning A, Rao A (1997) IKK-1 and IKK-2: cytokine-activated IkappaB kinases essential for NF-kappaB activation. Science 278: 860–6

Metzler M, Gan L, Mazarei G, Graham RK, Liu L, Bissada N, Lu G, Leavitt BR, Hayden MR (2010) Phosphorylation of huntingtin at Ser421 in YAC128 neurons is associated with protection of YAC128 neurons from NMDA-mediated excitotoxicity and is modulated by PP1 and PP2A. J Neurosci 30: 14318–29

Mishra R, Hoop CL, Kodali R, Sahoo B, van der Wel PC, Wetzel R (2012) Serine phosphorylation suppresses huntingtin amyloid accumulation by altering protein aggregation properties. J Mol Biol 424: 1–14

Neueder A, Landles C, Ghosh R, Howland D, Myers RH, Faull RLM, Tabrizi SJ, Bates GP (2017) The pathogenic exon 1 HTT protein is produced by incomplete splicing in Huntington’s disease patients. Sci Rep 7: 1307

Ochaba J, Fote G, Kachemov M, Thein S, Yeung SY, Lau AL, Hernandez S, Lim RG, Casale M, Neel MJ, Monuki ES, Reidling J, Housman DE, Thompson LM, Steffan JS (2019) IKKbeta slows Huntington’s disease progression in R6/1 mice. Proc Natl Acad Sci U S A 116: 10952–10961

Olshina MA, Angley LM, Ramdzan YM, Tang J, Bailey MF, Hill AF, Hatters DM (2010) Tracking mutant huntingtin aggregation kinetics in cells reveals three major populations that include an invariant oligomer pool. J Biol Chem 285: 21807–16

Perkins ND (2007) Integrating cell-signalling pathways with NF-kappaB and IKK function. Nat Rev Mol Cell Biol 8: 49–62

Polley S, Huang DB, Hauenstein AV, Fusco AJ, Zhong X, Vu D, Schrofelbauer B, Kim Y, Hoffmann A, Verma IM, Ghosh G, Huxford T (2013) A structural basis for IkappaB kinase 2 activation via oligomerization-dependent trans auto-phosphorylation. PLoS Biol 11: e1001581

Rangone H, Poizat G, Troncoso J, Ross CA, MacDonald ME, Saudou F, Humbert S (2004) The serum-and glucocorticoid-induced kinase SGK inhibits mutant huntingtin-induced toxicity by phosphorylating serine 421 of huntingtin. Eur J Neurosci 19: 273–9

Ratovitski T, O’Meally RN, Jiang M, Chaerkady R, Chighladze E, Stewart JC, Wang X, Arbez N, Roby E, Alexandris A, Duan W, Vijayvargia R, Seong IS, Lavery DJ, Cole RN, Ross CA (2017) Post-Translational Modifications (PTMs), Identified on Endogenous Huntingtin, Cluster within Proteolytic Domains between HEAT Repeats. J Proteome Res 16: 2692–2708

Ross CA, Tabrizi SJ (2011) Huntington’s disease: from molecular pathogenesis to clinical treatment. Lancet Neurol 10: 83–98

Sathasivam K, Neueder A, Gipson TA, Landles C, Benjamin AC, Bondulich MK, Smith DL, Faull RL, Roos RA, Howland D, Detloff PJ, Housman DE, Bates GP (2013) Aberrant splicing of HTT generates the pathogenic exon 1 protein in Huntington disease. Proc Natl Acad Sci U S A 110: 2366–70

Saudou F, Humbert S (2016) The Biology of Huntingtin. Neuron 89: 910–26

Schilling B, Gafni J, Torcassi C, Cong X, Row RH, LaFevre-Bernt MA, Cusack MP, Ratovitski T, Hirschhorn R, Ross CA, Gibson BW, Ellerby LM (2006) Huntingtin phosphorylation sites mapped by mass spectrometry. Modulation of cleavage and toxicity. J Biol Chem 281: 23686–97

Schrofelbauer B, Polley S, Behar M, Ghosh G, Hoffmann A (2012) NEMO ensures signaling specificity of the pleiotropic IKKbeta by directing its kinase activity toward IkappaBalpha. Mol Cell 47: 111–21

Sharma S, tenOever BR, Grandvaux N, Zhou GP, Lin R, Hiscott J (2003) Triggering the interferon antiviral response through an IKK-related pathway. Science 300: 1148–51

Simpson GL, Hughes JA, Washio Y, Bertrand SM (2009) Direct small-molecule kinase activation: Novel approaches for a new era of drug discovery. Curr Opin Drug Discov Devel 12: 585–96

Swingle M, Ni L, Honkanen RE (2007) Small-molecule inhibitors of ser/thr protein phosphatases: specificity, use and common forms of abuse. Methods Mol Biol 365: 23–38

Thompson LM, Aiken CT, Kaltenbach LS, Agrawal N, Illes K, Khoshnan A, Martinez-Vincente M, Arrasate M, O’Rourke JG, Khashwji H, Lukacsovich T, Zhu YZ, Lau AL, Massey A, Hayden MR, Zeitlin SO, Finkbeiner S, Green KN, LaFerla FM, Bates G et al. (2009) IKK phosphorylates Huntingtin and targets it for degradation by the proteasome and lysosome. J Cell Biol 187: 1083–99

Tsuchiya Y, Osaki K, Kanamoto M, Nakao Y, Takahashi E, Higuchi T, Kamata H (2017) Distinct B subunits of PP2A regulate the NF-kappaB signalling pathway through dephosphorylation of IKKbeta, IkappaBalpha and RelA. FEBS Lett 591: 4083–4094

Vezzoli E, Caron I, Talpo F, Besusso D, Conforti P, Battaglia E, Sogne E, Falqui A, Petricca L, Verani M, Martufi P, Caricasole A, Bresciani A, Cecchetti O, Rivetti di Val Cervo P, Sancini G, Riess O, Nguyen H, Seipold L, Saftig P et al. (2019) Inhibiting pathologically active ADAM10 rescues synaptic and cognitive decline in Huntington’s disease. J Clin Invest 129: 2390–2403

Weiss A, Trager U, Wild EJ, Grueninger S, Farmer R, Landles C, Scahill RI, Lahiri N, Haider S, Macdonald D, Frost C, Bates GP, Bilbe G, Kuhn R, Andre R, Tabrizi SJ (2012) Mutant huntingtin fragmentation in immune cells tracks Huntington’s disease progression. J Clin Invest 122: 3731–6

Weiss WA, Taylor SS, Shokat KM (2007) Recognizing and exploiting differences between RNAi and smallmolecule inhibitors. Nat Chem Biol 3: 739–44

William Yang X, Gray M (2011) Mouse Models for Validating Preclinical Candidates for Huntington’s Disease. In Neurobiology of Huntington’s Disease: Applications to Drug Discovery, Lo DC, Hughes RE (eds) Boca Raton (FL):

Xu X, Ng B, Sim B, Radulescu CI, Yusof N, Goh WI, Lin S, Lim JSY, Cha Y, Kusko R, Kay C, Ratovitski T, Ross C, Hayden MR, Wright G, Pouladi MA (2020) pS421 huntingtin modulates mitochondrial phenotypes and confers neuroprotection in an HD hiPSC model. Cell Death Dis 11: 809

Yu H, Lin L, Zhang Z, Zhang H, Hu H (2020) Targeting NF-kappaB pathway for the therapy of diseases: mechanism and clinical study. Signal Transduct Target Ther 5: 209

Yum S, Li M, Fang Y, Chen ZJ (2021) TBK1 recruitment to STING activates both IRF3 and NF-kappaB that mediate immune defense against tumors and viral infections. Proc Natl Acad Sci U S A 118

Zandi E, Rothwarf DM, Delhase M, Hayakawa M, Karin M (1997) The IkappaB kinase complex (IKK) contains two kinase subunits, IKKalpha and IKKbeta, necessary for IkappaB phosphorylation and NF-kappaB activation. Cell 91: 243–52

Zheng Z, Li A, Holmes BB, Marasa JC, Diamond MI (2013) An N-terminal nuclear export signal regulates trafficking and aggregation of Huntingtin (Htt) protein exon 1. J Biol Chem 288: 6063–71

